# Temporal structure of task engagement reorganizes infra-slow BOLD dynamics

**DOI:** 10.64898/2026.05.27.727996

**Authors:** Yujia Ao, Mingjun Chi, Yasir Catal, Zirui Huang, Chao Wang, Xue Mei Song, Zixiao Liu, Xi-Nian Zuo, Yifeng Wang, Georg Northoff

## Abstract

Task-based fMRI has traditionally characterized cognitive engagement through spatial patterns of BOLD activation, leaving open whether tasks also organize activity along the frequency dimension. Here we tested whether temporally structured task engagement reshapes infra-slow BOLD dynamics, asking whether task responses are simply superimposed onto ongoing activity or instead redistribute power across the infra-slow spectrum. Across three independent fMRI datasets, task timing systematically reorganized the BOLD power spectrum. In the main visual attention dataset, periodic stimulation produced sharp stimulus-locked peaks at 0.083 and 0.125 Hz; critically, these gains were matched by a reduction concentrated within the dominant intrinsic infra-slow range (0.01-0.04 Hz), indicating redistribution rather than a simple local increase. Independent auditory and visual-motion datasets showed analogous task-frequency responses at their respective timescales, indicating that this reorganization generalizes across modalities and stimulation frequencies. The peaks were spatially related to GLM-derived activation but persisted after removal of the GLM-modeled response, indicating that this frequency-specific organization is not fully reducible to conventional task-evoked activation. These features were functionally informative: task-imposed frequencies distinguished cognitive states, predicted individual reaction time, and preserved subject-specific signatures across runs more effectively than broader spectral ranges or GLM-derived features. Finally, Jansen-Rit simulations coupled to a Balloon-Windkessel model reproduced the peaks, showing that their expression depends on balanced excitatory, inhibitory, and filter-gain regimes rather than simple amplification of input. Together, these findings indicate that task-fMRI reveals not only where cognition is localized, but how temporally structured engagement redistributes infra-slow dynamics across timescales.

## Introduction

Task-based fMRI has traditionally characterized cognitive engagement through regional changes in BOLD amplitude — task-evoked increases and decreases that are typically modeled as responses superimposed onto an ongoing baseline of spontaneous activity. This framework, most commonly formalized through the general linear model (GLM), has provided a powerful approach for mapping task-evoked activation and deactivation across cortical systems (Friston et al., 1994; Shulman et al., 1997; Zhang et al., 2023). Yet cognition unfolds not only across cortical space but across a hierarchy of temporal scales (Buzsáki, 2025; Golesorkhi et al., 2021a; Gong and Zuo, 2025; Palva and Palva, 2018; Wolff et al., 2022); it remains open whether task engagement merely adds local responses to this spontaneous baseline in a task-specific frequency or timescales, or instead globally reorganizes the brain’s ongoing dynamics along the whole hierarchy of its timescales (Sasai et al., 2021). This question is particularly relevant for infra-slow BOLD fluctuations, which constitute a prominent component of large-scale fMRI dynamics and have been linked to functional network organization, intrinsic timescales, and behavioral variability (Bola and Sabel, 2015; Golesorkhi et al., 2021b; Gutierrez-Barragan et al., 2019; Waschke et al., 2021). At faster electrophysiological timescales, steady-state evoked potential (SSEP), neural entrainment, and frequency-tagging approaches have shown that rhythmic sensory input can elicit neural responses at the stimulation frequency and its harmonics, providing a sensitive marker of sensory processing, attention, and cognitive state (Lakatos et al., 2019; Lu et al., 2017; Norcia et al., 2015; Vialatte et al., 2010; Wieser et al., 2016). Whether this frequency-tagging logic extends to fMRI and its infra-slow timescales remains less clear.

Recent work suggests that slow periodic stimulation can induce low-frequency steady-state BOLD responses (lfSSBR), with spectral peaks emerging at the task-imposed frequency (Kasagi et al., 2017; Klar et al., 2023a, 2023b; Wang et al., 2018; Wolman et al., 2024). These studies established that lfSSBR exists, but characterized it largely through the peak itself, leaving open whether it is an isolated response or the local signature of a broader, more global, spectrum-wide redistribution in which the temporal scales imposed by the task manifest within the brain’s own frequency architecture. This distinction yields two divergent predictions. If task responses are simply superimposed onto ongoing activity, power should rise at the task-imposed frequencies while the rest of the spectrum — including the brain’s dominant intrinsic low-frequency fluctuations — remains essentially preserved (Fox et al., 2006; He, 2013; Huang et al., 2017). The resulting spectral change would therefore amount primarily to the frequency-domain footprint of the task-evoked response already captured by a conventional GLM.

Alternatively, if task engagement actively reorganizes ongoing dynamics, the same task-related temporal structure should redistribute power across the infra-slow spectrum as a whole, such that gains at the task-imposed frequencies are matched by a frequency-specific reduction concentrated within the dominant intrinsic infra-slow range; this is, consistent with previous evidence that task performance suppresses spontaneous low-frequency BOLD power, signal variability, and scale-free dynamics (Bianciardi et al., 2009; Churchill et al., 2016; Duff et al., 2008; He, 2011). On this view, the local spectral peak is not a response added onto an otherwise preserved spectrum but the signature of a global reallocation of infra-slow power across timescales, and such redistribution should remain detectable after the GLM-modeled response is removed, as suggested by prior observations of altered task-residual low-frequency activity following regression of event-related signals (Zhang and Li, 2012). The decisive question, then, is not whether a spectral peak appears, but whether that gain is reducible to conventional activation or instead reflects a spectrum-wide redistribution that activation analyses alone cannot capture.

If this global redistribution reflects a general principle of how temporally structured engagement reorganizes infra-slow dynamics, it should generalize across sensory modalities and stimulation timescales rather than being tied to any single task or a particular modality or domain. Because such task-general demands (Nau et al., 2024) unfold over the same seconds-to-tens-of-seconds timescale as infra-slow BOLD dynamics (Palva and Palva, 2012; Weissman et al., 2006). If such global spectral reorganization is functionally relevant rather than incidental, its spectral features should further distinguish cognitive states, predict behavioral efficiency, and identify individuals more effectively than conventional activation-based features.

Establishing frequency-specific organization of infra-slow BOLD dynamics, however, leaves a central mechanistic ambiguity: does lfSSBR simply reflect stronger periodic input, or does it arise because cognitive engagement shifts a gain regime through which all ongoing activity is filtered? Attentional prioritization is widely thought to depend on gain control (Reynolds and Heeger, 2009) — the regulation of cortical excitability that amplifies task-relevant inputs relative to competing intrinsic activity (Harris and Thiele, 2011). Such gain control is shaped by neuromodulatory influences, including cholinergic and noradrenergic systems, which have been linked to sensory excitatory/inhibitory gain, attentional selectivity, arousal, and temporal filtering (Coronel-Oliveros et al., 2021; Thiele and Bellgrove, 2018). Although these influences cannot be isolated directly with conventional fMRI, biophysically grounded neural mass models provide a principled way to test whether gain-like changes in excitation, inhibition, and filtering are sufficient to redistribute infra-slow power into the observed frequency-specific organization, that is, whether lfSSBR reflects a regulated, balanced gain regime rather than simple amplification of external input (Moran et al., 2007).

Here, across three independent fMRI datasets and a biophysical neural mass model, we asked whether task engagement with temporally structured events reorganizes the whole infra-slow power spectrum, rather than simply adding local responses to it (Fig. 1). We first examined whether task timing reshapes this spectrum, concentrating power into narrowband stimulus-locked peaks (lfSSBR) under regular rhythmic stimulation and producing broader redistribution across task-relevant frequencies under less regular timing; in both cases, gains at task-imposed frequencies were matched by losses in the dominant intrinsic infra-slow range. We then asked whether this reorganization is reducible to conventional task-evoked activation, whether its spectral features carry information about cognitive state, behavioral efficiency, and individual identity, and whether it can be reproduced by a neural mass model through gain-dependent filtering rather than simple amplification of external drive. Together, these analyses test the idea that task-related BOLD activity reflects not only where cognition modulates regional activation, but also how temporally structured engagement reorganizes the brain’s whole infra-slow power spectrum — of which the local stimulus-locked peak is just one expression.

**Figure 1.**
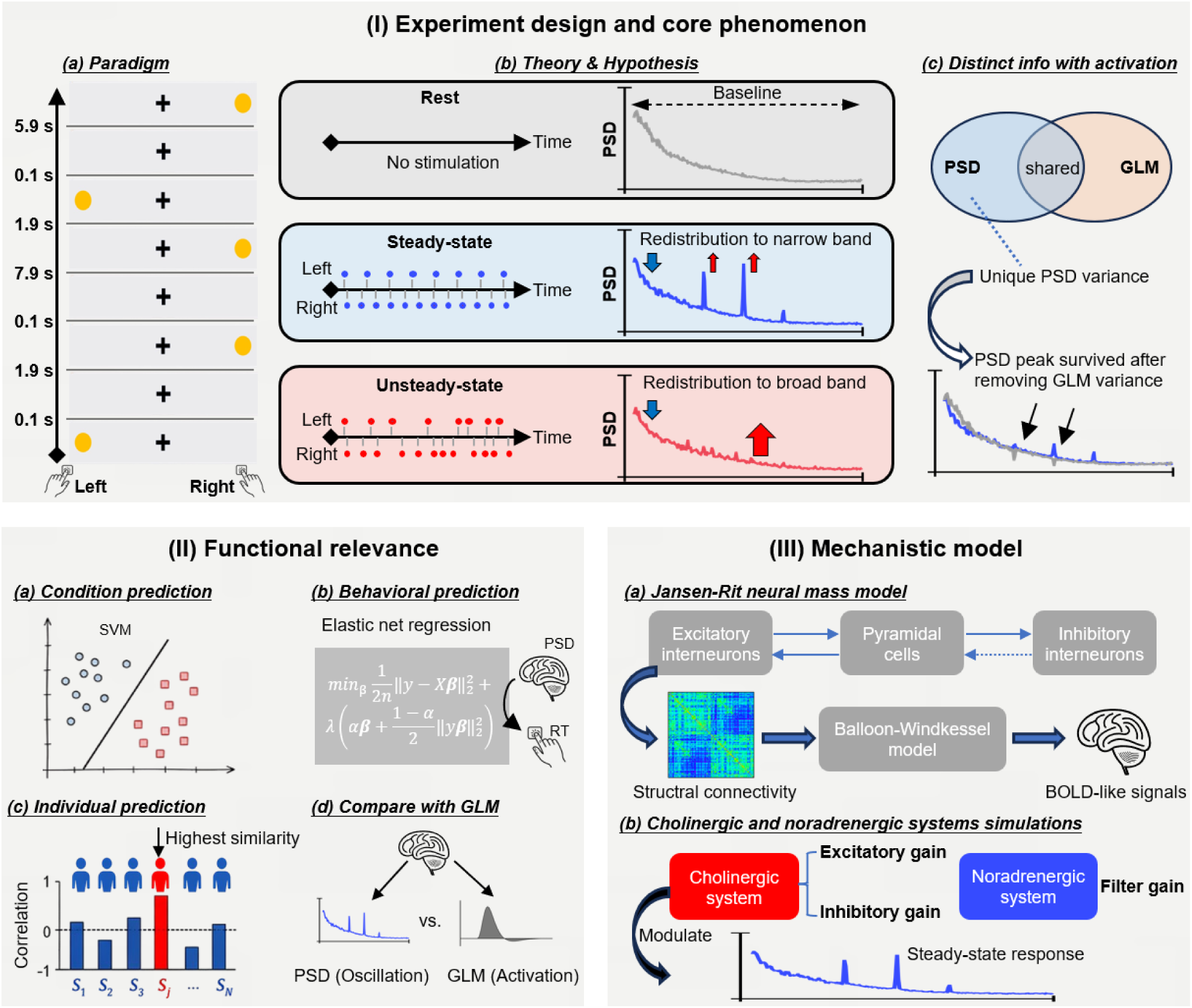
Overview of the study design, core phenomenon, functional relevance, and mechanistic model. (I) Experiment design and core phenomenon. (a) Experimental paradigm: left and right visual targets were presented at the times indicated, requiring left/right button-press responses. (b) Theory and hypothesis. Three conditions with distinct temporal structures: resting state (no stimulation), steady-state (periodic left-right stimulation at fixed intervals), and unsteady-state (aperiodic stimulation), together with their hypothesized BOLD power spectra (PSD). Relative to rest, both task conditions are predicted to redistribute power away from the dominant low infra-slow range (blue arrow) and toward task-imposed frequencies: steady-state stimulation concentrates power into narrowband stimulus-locked peaks (lfSSBR; red arrows), whereas unsteady-state stimulation produces a broader redistribution across task-relevant frequencies (red arrow). (c) PSD carries information that is distinct from, though partly shared with, GLM-derived activation: stimulus-locked PSD peaks persist after regressing out the GLM-modeled response, reflecting a component of unique PSD variance. (II) Functional relevance. (a) Support vector machine (SVM) classification of the three conditions from spectral features. (b) Elastic net regression predicting individual reaction time (RT) from PSD features. (c) Subject-level identification based on spectral profiles. (d) Comparison between PSD-based and GLM-based features, where applicable (behavioral prediction and individual identification). (III) Mechanistic model. (a) A Jansen–Rit neural mass model (excitatory interneurons, pyramidal cells, inhibitory interneurons) coupled via empirical structural connectivity and convolved with the Balloon–Windkessel model to generate BOLD-like signals. (b) Simulated cholinergic (excitatory/inhibitory gain) and noradrenergic (filter gain) modulation of the steady-state spectral response.

## Results

### Task-imposed spectral redistribution of infra-slow BOLD power

We first asked whether task engagement reorganized the distribution of BOLD power across infra-slow frequencies in the main UESTC dataset. For each condition, we computed the fractional power spectral density (PSD) of regional BOLD signals and averaged power across the whole brain. This normalization of PSD was applied to isolate changes in spectral shape from condition-related differences in overall BOLD amplitude, thereby directly characterizing how power was redistributed across frequencies independent of global scaling effects. The most prominent low-frequency steady-state BOLD responses (lfSSBR) were observed in the steady-state condition, where sharp peaks emerged at the stimulus-locked frequencies of 0.083 Hz and 0.125 Hz (Fig. 2A). These peaks were absent or markedly attenuated during rest and the unsteady-state condition, as confirmed by paired-sample t-tests. Spatially, the corresponding effects were broadly distributed across the cortex rather than being restricted to the directly stimulated sensory and motor systems (Fig. 2A and Supplementary Fig. 1).

**Figure 2.**
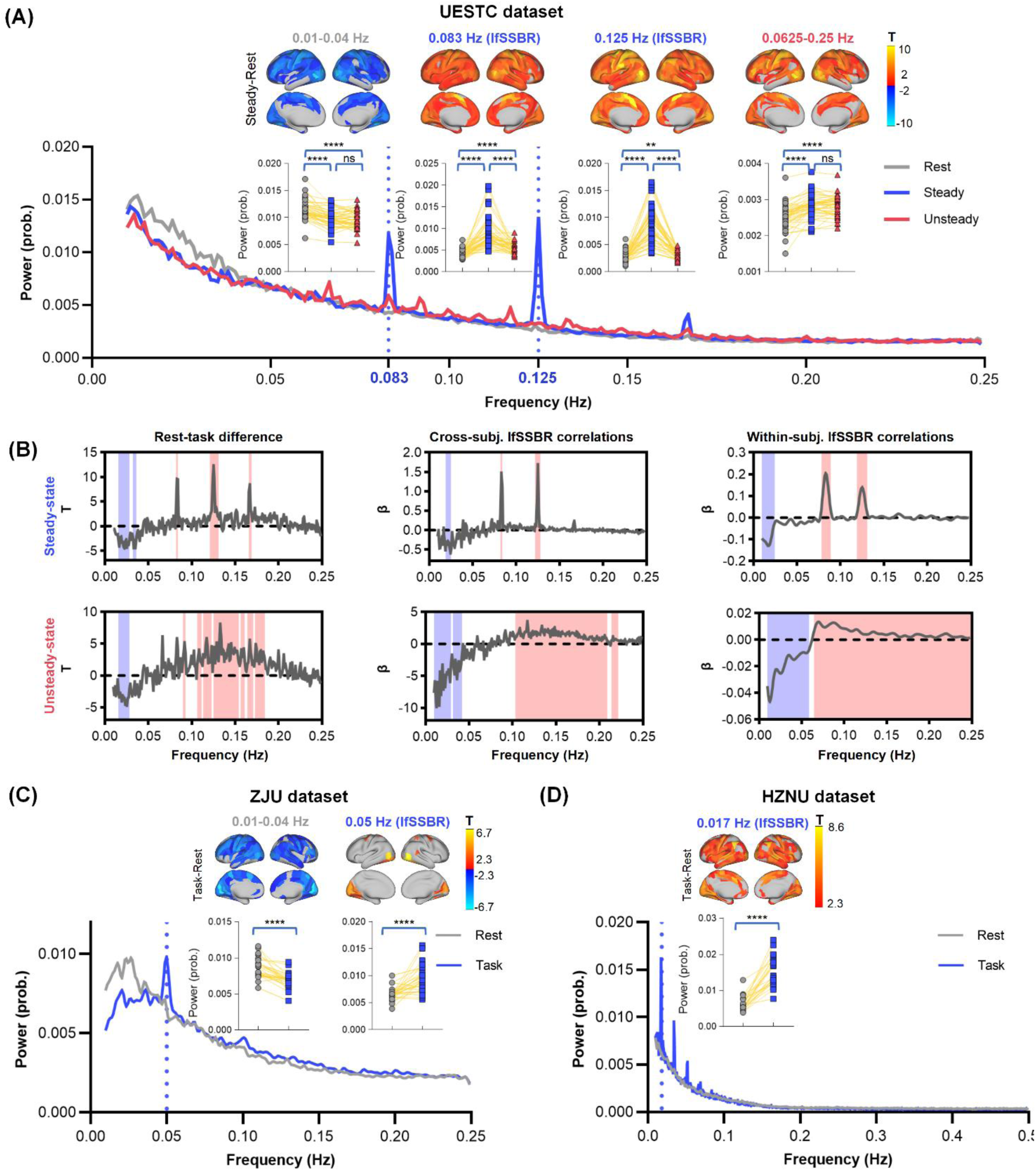
Infra-slow power redistribution across conditions and datasets. (A) UESTC dataset. Whole-brain averaged BOLD power spectral density (PSD) is shown for rest, steady-state, and unsteady-state conditions. Steady-state stimulation produced low-frequency steady-state BOLD responses (lfSSBR) at 0.083 Hz and 0.125 Hz. Colored bars indicate the frequency ranges of interest: 0.01-0.04 Hz, the stimulus-locked frequencies of 0.083 Hz and 0.125 Hz, and the broader task-relevant range of 0.0625-0.25 Hz. Cortical maps show vertex-wise t-statistics for the differences between steady-state condition and resting-state condition (gray regions are non-significant, see Supplementary Fig. 1 showing results for other condition pairs), and insets show paired subject-level comparisons. (B) Frequency-resolved analyses for the steady-state (top) and unsteady-state (bottom) conditions. Left, rest-task difference (paired t-test) across the spectrum; Middle, cross-subject association between lfSSBR power and power at every other frequency (y-axis, regression slope β). Right, the mean within-subject association between lfSSBR power and every other frequency across time series, estimated with a sliding-window approach. Multiple comparisons across frequency bins were controlled using one-dimensional cluster-based permutation testing. (C) ZJU dataset. Task engagement reduced low infra-slow power at 0.01-0.04 Hz and increased power at the block-design frequency of 0.05 Hz. (D) HZNU dataset. Task engagement produced a spectral peak at approximately 0.017 Hz relative to rest, consistent with the long inter-trial intervals of the auditory task. Asterisks indicate statistical significance (\**p* < 0.05, \*\**p* < 0.01, \*\*\**p* < 0.001, \*\*\*\**p* < 0.0001; ns, not significant). Cortical maps and paired tests were corrected for multiple comparisons using FDR.

We next examined whether these lfSSBR were accompanied by broader changes in the spectral profile. During rest, BOLD power was concentrated in the low infra-slow range of 0.01-0.04 Hz, whereas both task conditions showed reduced power in this band. Conversely, power in the broader task-relevant range of 0.0625-0.25 Hz was elevated during task conditions relative to rest, with a pronounced increase also observed in the unsteady-state condition (Fig. 2A). Thus, task engagement did not produce a uniform increase or decrease in BOLD power. Instead, it redistributed power away from dominant spontaneous low-frequency fluctuations and toward frequencies aligned with the temporal structure of the task.

To localize this redistribution without predefining frequency bands, we further adopted a data-driven, frequency-resolved approach. Comparing task and rest frequency by frequency, the only significant task-related decrease fell in the very slow band corresponding to our predefinition of 0.01-0.04 Hz, whereas the stimulus-locked frequencies and their harmonics showed significant increases, in both the steady- and unsteady-state conditions (Fig. 2B, left). We then asked which frequencies covaried with lfSSBR power directly. Across subjects, lfSSBR power was negatively associated with power exclusively in the 0.01-0.04 Hz band, with no comparable association at any other frequency (Fig. 2B, middle); the same selectivity held within subjects when lfSSBR power was tracked across sliding windows (Fig. 2B, right). Together, these analyses localize the trade-off to a specific exchange between slow infra-slow fluctuations and lfSSBR, indicating a genuine redistribution of power from slow to stimulus-locked frequencies.

As a complementary check using absolute rather than relative power, we repeated the same analyses on unnormalized PSD in the main UESTC dataset (Supplementary Fig. 2A). Raw PSD showed an overall reduction in BOLD power during task conditions relative to rest, consistent with task-related suppression of spontaneous BOLD fluctuations (Ao et al., 2025; Waschke et al., 2021). Importantly, the key spectral effects were preserved: power remained elevated at the stimulus-locked frequencies of 0.083 Hz and 0.125 Hz, whereas power in the low infra-slow range of 0.01-0.04 Hz was reduced. These findings indicate that the observed redistribution was not an artifact of total-power normalization or global spectral scaling. To further exclude motion-related confounds, we computed the PSD of mean framewise displacement. No significant condition differences were observed in the four frequency ranges of interest, namely 0.083 Hz, 0.125 Hz, 0.01-0.04 Hz, and 0.0625-0.25 Hz, suggesting that the spectral redistribution was not explained by head motion (Supplementary Fig. 2B).

We then tested whether this temporal organization generalized across independent datasets with different task structures. In the HZNU dataset, an auditory task imposed a much slower temporal rhythm, with inter-trial intervals of 52-60 s (approximately 0.017-0.019 Hz). Consistent with this timing, task engagement produced a clear spectral peak at 0.017 Hz relative to rest, accompanied by widespread task-related increases in the corresponding cortical map (Fig. 2C). In the ZJU dataset, which used a visual motion-detection task with a block-design frequency of 0.05 Hz, task engagement reduced power in the low infra-slow range of 0.01-0.04 Hz while increasing power at 0.05 Hz, again matching the temporal structure of the task (Fig. 2D).

Together, these results show that task-related BOLD power changes are organized by the temporal structure of stimulation. Across three independent datasets, task engagement reduced power in spontaneous low-frequency ranges and enhanced power at task-imposed frequencies, whether these frequencies arose from distinct sensory modalities, rhythmic stimulation, or block-design timing.

### lfSSBR is not fully explained by GLM-derived activation

We next asked whether the stimulus-locked spectral peaks could be explained by conventional task-evoked BOLD activation. GLM analyses revealed robust task-related activation patterns, including positive responses in contralateral visual and sensorimotor cortices and characteristic deactivations within the default mode network (DMN; Fig. 3A).

**Figure 3.**
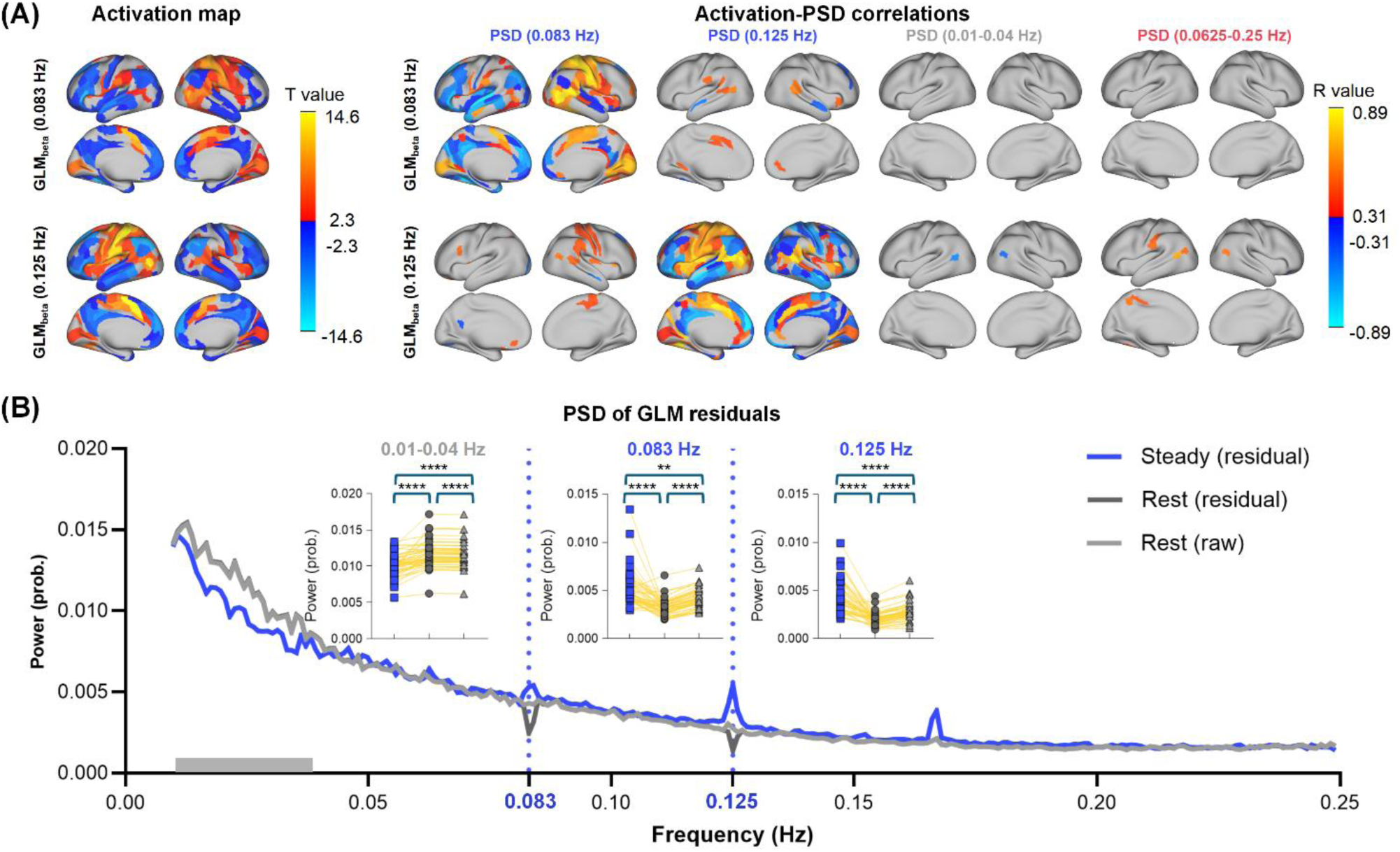
lfSSBR persists after removal of GLM-modeled activation. (A) Left: Cortical t-statistic maps of GLM-derived activation in the steady-state condition for the two stimulus-locked frequencies, 0.083 Hz and 0.125 Hz. Right: For each cortical parcel, Pearson correlations were computed across subjects between GLM beta values and PSD, and the resulting correlation coefficients were displayed on the cortical surface. Rows indicate GLM beta maps for the 0.083-Hz and 0.125-Hz regressors; columns indicate PSD at 0.083 Hz, 0.125 Hz, 0.01-0.04 Hz, and 0.0625- 0.25 Hz. (B) Whole-brain averaged PSD of steady-state GLM residuals, compared with raw resting-state PSD and procedure-matched resting-state GLM residuals. Peaks at 0.083 Hz and 0.125 Hz persisted after GLM regression, indicating that lfSSBR was not fully reducible to canonical task-evoked activation. Insets show paired subject-level comparisons.

We then examined the relationship between spectral power and GLM-derived activation across frequencies. This analysis revealed a frequency-specific coupling that was selectively expressed in the steady-state condition. Power at the stimulus-locked frequencies of 0.083 Hz and 0.125 Hz was positively correlated with GLM beta values in task-positive regions, but negatively correlated in the DMN (Fig. 3A). Thus, stronger stimulus-locked power was associated with both greater task-positive recruitment and stronger DMN suppression. In contrast, correlations at non-stimulus-locked frequencies were weak or absent, and no comparable relationships were observed in the unsteady-state condition (Supplementary Fig. 3A). In the independent HZNU dataset (auditory steady-state task at 0.017 Hz), GLM analyses revealed strong activation in auditory cortices; however, activation-PSD correlations were sparser and did not co-localize with the activation map (Supplementary Fig. 4A), suggesting that spectral power carries information beyond canonical task-evoked activation. Together, these findings indicate that task-locked spectral power tracks, but is not reducible to, the magnitude of task-evoked activation and deactivation in a frequency-specific manner.

To determine whether these spectral peaks were fully accounted for by canonical GLM-modeled responses, we recomputed the PSD after regressing out the GLM-predicted response from the steady-state BOLD signals. Because the GLM regressors shared temporal structure with the stimulus-locked stimulation, this procedure could in principle remove not only conventional task-evoked activation but also oscillatory power at the same frequencies. We therefore compared residual steady-state power with two resting-state baselines: raw resting-state power and resting-state power after applying the same GLM regression procedure. This analysis for resting-state power tested whether the residual spectral feature patterns could arise from spurious fitting of task regressors to ongoing BOLD fluctuations.

lfSSBR at 0.083 Hz and 0.125 Hz remained clearly visible in the GLM residuals and were significantly higher than both resting-state baselines (Fig. 3B). Reduced power in the low infra-slow range was also preserved after GLM regression. A similar residual redistribution pattern was observed in the unsteady-state condition, with reduced low infra-slow power and elevated broadband task-related power after GLM regression (Supplementary Fig. 3B). The 0.017 Hz peak in the HZNU dataset likewise persisted after GLM regression (Supplementary Fig. 4B).

Together, these findings indicate that lfSSBR overlaps with conventional task-evoked activation but is not reducible to it. Instead, it captures a residual frequency-specific component of task-related BOLD dynamics that persists after removal of the canonical GLM-modeled response. More broadly, the persistence of spectral redistribution in both steady-state and unsteady-state residuals suggests that task-related infra-slow activity carries temporal structure that is not fully captured by conventional activation amplitude.

### lfSSBR predicts cognitive state, behavior, and individual identity

We next tested whether the frequency-specific spectral features identified above carried functionally meaningful information. We evaluated this at three levels: discrimination of cognitive state, prediction of behavioral performance, and identification of individual subjects across runs. Where applicable, we compared PSD-based features with conventional GLM-derived activation measures.

First, we asked whether spectral power could distinguish resting, steady-state, and unsteady-state conditions in the main UESTC dataset. We trained support vector machine (SVM) classifiers with leave-one-out cross-validation using PSD features (ROIs * Subjects) from different frequency ranges. PSD features at the stimulus-locked frequencies of 0.083 Hz and 0.125 Hz achieved significant three-way state classification (macro-average AUC = 0.88 and 0.93, both permutation *p < 0.0001*), with each frequency preferentially discriminating the condition most strongly associated with its temporal structure (Fig. 4A). By contrast, broad frequency ranges showed substantially weaker discrimination: power in the low infra-slow range of 0.01-0.04 Hz yielded significant AUC = 0.63 (permutation *p < 0.0001*), and power in the broader task-relevant range of 0.0625-0.25 Hz yielded significant AUC = 0.68 (permutation *p < 0.0001*), both significantly below the stimulus-locked features (all pairwise DeLong tests, FDR-corrected *p* < 0.01). Per-class confusion matrices confirmed that broad-band features were particularly poor at discriminating task conditions, with per-class accuracy falling near or below chance for steady-state and unsteady-state (Supplementary Fig. 5A). GLM-derived features were not included in this analysis because GLM regressors are defined from prior knowledge of the experimental design, making them unsuitable as an independent benchmark for classifying the same task conditions. To test generalizability, we repeated the classification in the HZNU and ZJU datasets using their respective task-locked frequencies; performance remained substantially above chance in both datasets (Supplementary Fig. 6).

**Figure 4.**
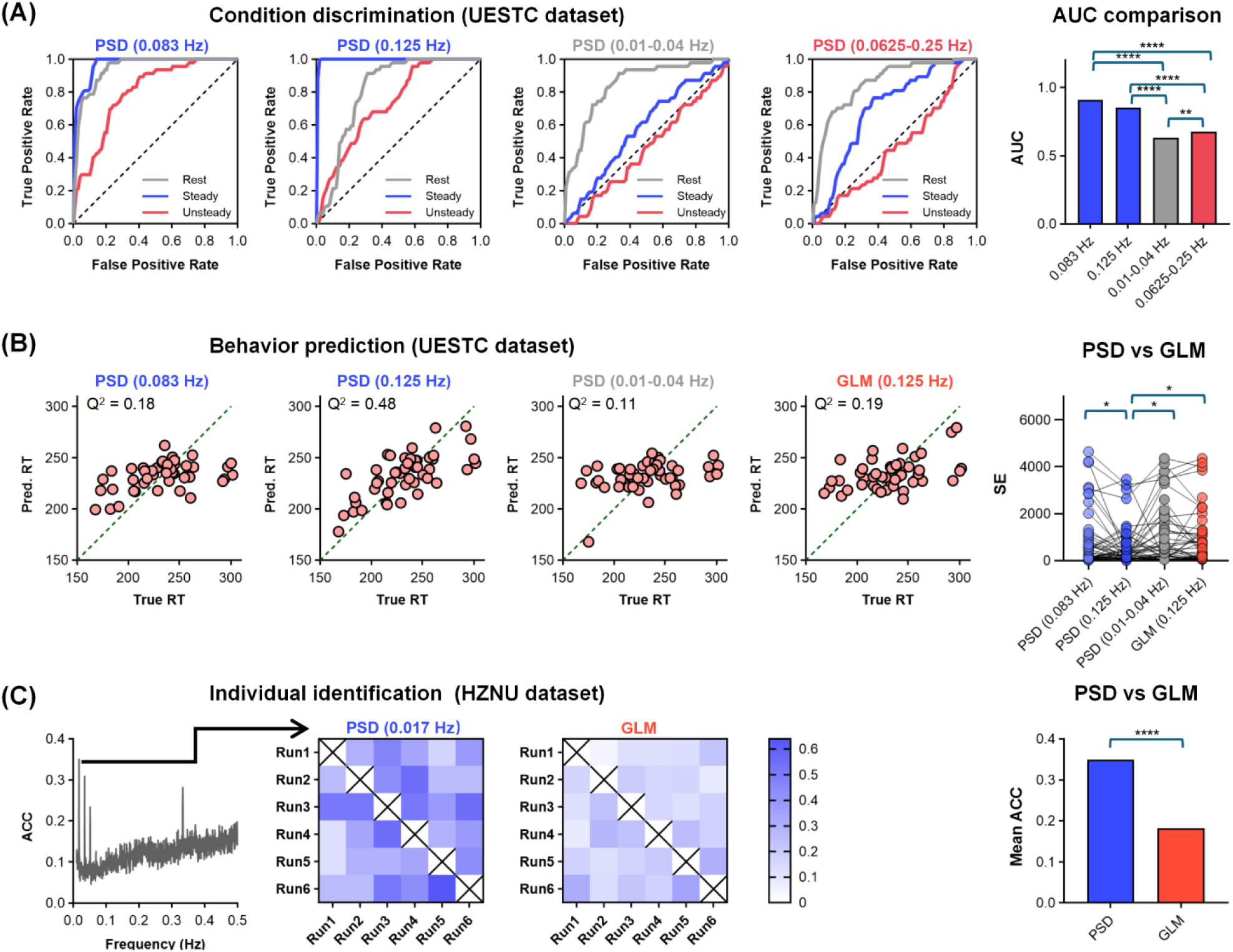
lfSSBR predicts cognitive states, behavior, and individual identity. (A) Condition prediction (UESTC dataset). Left: Receiver operating characteristic (ROC) curves for three-class SVM classification (rest, steady-state, unsteady-state) using spectral power as input features, computed separately for four frequency bands: stimulus-locked frequencies (0.083 Hz, 0.125 Hz), the infra-slow band (0.01-0.04 Hz), and the broadband task-relevant range (0.0625-0.25 Hz). Classification used a one-versus-rest scheme with leave-one-out cross-validation. Right: Area under the curve (AUC) compared across frequency bands for each frequency. Pairwise AUC comparisons used DeLong’s test. (B) Behavior prediction (UESTC dataset). Left: Scatter plots of predicted versus true reaction time (RT) from elastic net regression using four feature sets: PSD at 0.01-0.04 Hz, 0.083 Hz, 0.125 Hz, and GLM beta at 0.125 Hz. Predictive performance is quantified by Q². Right: Subject-wise squared error (SE) compared across feature sets. (C) Individual prediction (HZNU dataset). Left: Identification accuracy (ACC) across frequencies, peaking in the 0.017 Hz range corresponding to the stimulus-locked band. Middle: Run-by-run cross-correlation matrices of subject-level spectral profiles (PSD, 0.017 Hz) and GLM beta maps across six task runs; higher on-diagonal values indicate stronger within-subject identifiability. Right: Mean identification accuracy compared between PSD- and GLM-based features.

We next tested whether stimulus-locked spectral features carried behaviorally relevant information. Given the ceiling effects observed for accuracy (Supplementary Fig. 7), we focused the predictive analysis on reaction time (RT). Because PSD features consisted of many spatially distributed and correlated regional predictors relative to the sample size, regularized regression was required to limit overfitting and obtain stable out-of-sample predictions (Zou and Hastie, 2005). We therefore used elastic net regression, which combines L1- and L2-regularization, allowing sparse feature weighting while retaining groups of correlated predictors that may jointly carry behavioral information (Abram et al., 2016; Grosenick et al., 2013). In the UESTC dataset, elastic net models were trained to predict individual RT from PSD features and were benchmarked against models using GLM-derived activation features. In steady state, PSD at the 0.125 Hz stimulation frequency showed the strongest predictive performance, reaching a cross-validated Q² of 0.48 (permutation *p < 0.001*) and outperforming PSD at 0.083 Hz, low-frequency PSD in the 0.01-0.04 Hz range, and GLM beta estimates at the stimulation frequencies (Fig. 4B). Importantly, this advantage was robust across the elastic net lasso-ridge mixing parameter: although absolute Q² values varied with the degree of sparsity, the 0.125 Hz PSD model consistently showed the best or near-best performance across the parameter range (Supplementary Fig. 5B). During unsteady state, no significant Q² values survived after correction for multiple comparisons. These results indicate that individual differences in behavior were most strongly captured by stimulus-locked spectral power, especially at 0.125 Hz, rather than by conventional GLM activation or nonspecific low-frequency power.

Finally, we examined whether stimulus-locked spectral features preserved subject-specific information across repeated task runs in the HZNU dataset. Individual identification accuracy peaked at 0.017 Hz, matching the task-imposed temporal frequency (Fig. 4C). Cross-run similarity matrices showed stronger within-subject consistency for PSD-based features than for GLM-based features, and mean identification accuracy was significantly higher for PSD than for GLM. These results indicate that task-imposed spectral power not only reflects the temporal structure of stimulation but also preserves stable individual signatures across runs.

Together, these findings show that frequency-specific spectral power, especially at task-imposed frequencies, is functionally informative across multiple levels. It distinguishes cognitive states, predicts behavioral performance, and preserves individual identity more effectively than broader spectral ranges or, where applicable, conventional GLM-derived activation. These results extend the observed spectral redistribution from a descriptive frequency-domain phenomenon to a functionally relevant marker of task-related brain dynamics.

### Jansen-Rit modeling reveals steady-state-specific tuning of lfSSBR

To test whether the observed spectral redistribution could be reproduced by a biophysically motivated neural mass model, we implemented a Jansen-Rit model coupled with a Balloon-Windkessel hemodynamic model to generate BOLD-like signals (Fig. 5A). The model received either temporally regular input at 0.125 Hz or temporally irregular input with event intervals spanning the 0.0625-0.25 Hz range. In a single-node implementation, the simulated BOLD signals reproduced the main empirical pattern (Supplementary Fig. 8): steady-state input produced a pronounced lfSSBR at 0.125 Hz, whereas unsteady-state input did not generate a comparable narrowband peak but instead showed broader power redistribution across the stimulated frequency range. Resting-state activity, in contrast, remained dominated by low infra-slow power. Thus, the model captured the key descriptive distinction between temporally regular and temporally irregular task input.

**Figure 5.**
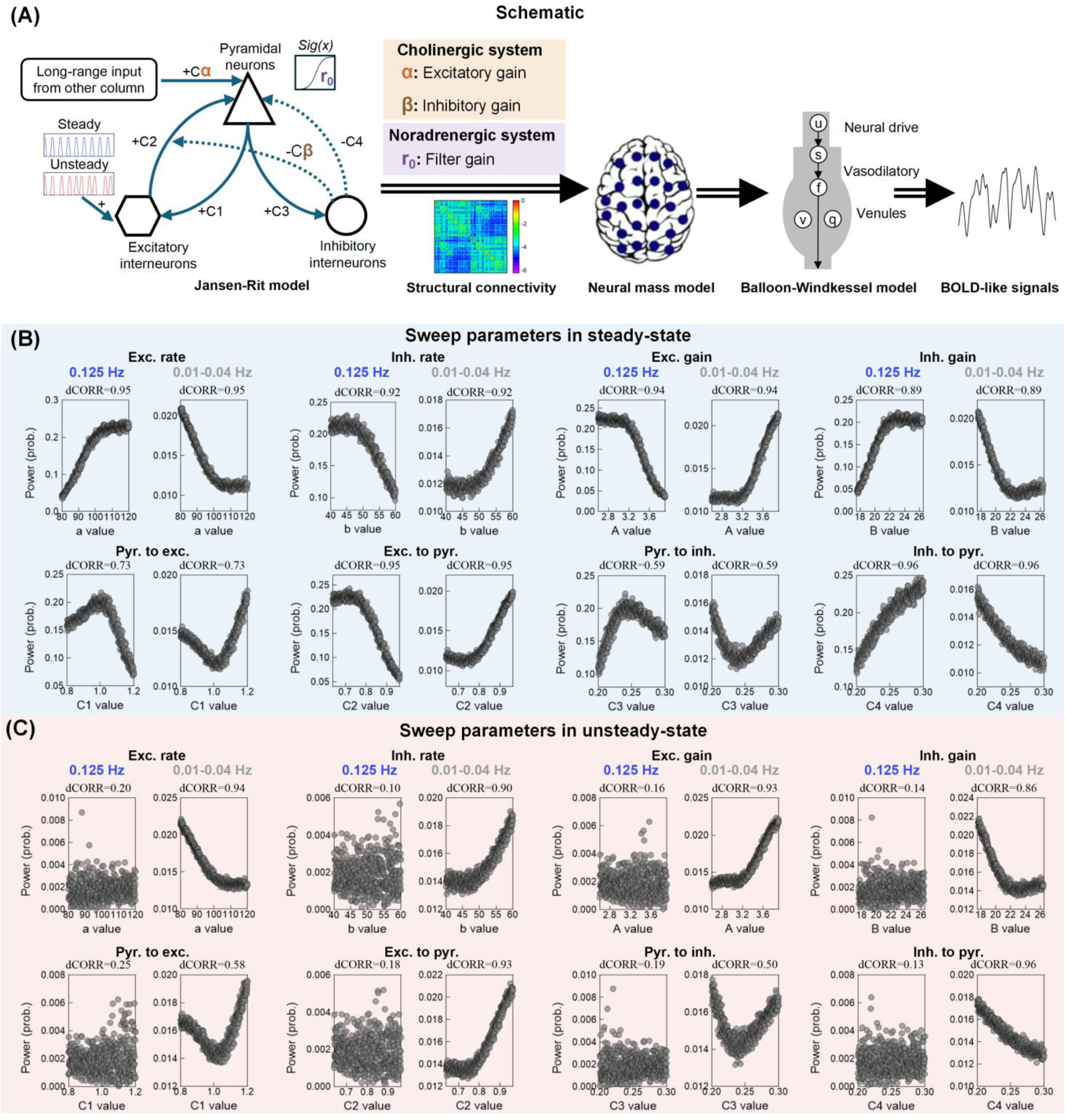
Jansen-Rit neural mass model reproduces lfSSBR. (A) Modeling pipeline. The local Jansen-Rit circuit comprises pyramidal cells (triangle), excitatory interneurons (hexagon), and inhibitory interneurons (circle), connected via four intrinsic coupling constants (*C1*: pyramidal to excitatory; *C2*: excitatory to pyramidal; *C3*: pyramidal to inhibitory; *C4*: inhibitory to pyramidal). excitatory and inhibitory synaptic rate constants are governed by *a* and *b*; excitatory and inhibitory synaptic gains by *A* and *B* (see Methods for detail). The cholinergic system modulates excitatory *α* and inhibitory *β* gains, and the noradrenergic system modulates filter gain *r₀*. Local circuits are coupled via empirical structural connectivity to form a whole-brain neural mass model, and the simulated neural activity was transformed into BOLD-like signals using the Balloon-Windkessel hemodynamic model. (B) Parameter sweeps under steady-state stimulation in single node. For each of the eight Jansen-Rit parameters (top row: EPSP *a*, IPSP *b*, excitatory gain *A*, inhibitory gain *B*; bottom row: coupling constants *C1*-*C4*), simulated whole-brain power is plotted against parameter value at the stimulus-locked frequency (0.125 Hz; blue) and the infra-slow band (0.01-0.04 Hz; gray). Distance correlation (dCORR) quantifying the parameter-power dependence is reported above each subplot. (C) Same parameter sweeps under unsteady-state stimulation. Infra-slow power (0.01-0.04 Hz) retains strong parameter dependence (dCORR = 0.50-0.96), whereas power at 0.125 Hz shows weak dependence (dCORR = 0.10-0.25).

We then asked whether this distinction depended on local circuit parameters. We systematically varied eight parameters governing synaptic dynamics, neuronal gains, and recurrent coupling strengths, and quantified simulated power at 0.125 Hz and in the low infra-slow range of 0.01-0.04 Hz. Under steady-state stimulation, 0.125 Hz power showed structured parameter dependence across local circuit parameters (distance correlation, dCORR = 0.59-0.96; Fig. 5B). Critically, low infra-slow power (0.01-0.04 Hz) showed the opposite pattern across the same parameter space, indicating a redistribution of spectral power from intrinsic low-frequency fluctuations toward the imposed stimulation frequency.

By contrast, under unsteady-state stimulation, this parameter-sensitive organization was much weaker at 0.125 Hz. Although low infra-slow power remained strongly modulated by local circuit parameters (dCORR = 0.50-0.96), power at 0.125 Hz showed only weak parameter dependence and lacked a coherent profile (dCORR = 0.10-0.25; Fig. 5C). These results indicate that local Jansen-Rit dynamics can reproduce the empirical distinction between steady-state and unsteady-state stimulation: temporally regular input produces a parameter-sensitive concentration of power at the imposed frequency, whereas temporally irregular input produces broader spectral redistribution without an equivalent narrowband peak.

Together, these simulations reproduced the empirical spectral findings in a biophysically motivated model and established that task/stimulus-frequency-specific power concentration can emerge from local neural mass dynamics. We next extended this framework to the whole-brain level to examine how cholinergic and noradrenergic gain parameters shaped the propagation of stimulus-locked spectral power changes across cortical networks.

### Neuromodulatory gain shapes lfSSBR

To examine how neuromodulatory gain parameters shape the network expression of lfSSBR, we delivered 0.125-Hz input to visual (VIS) and somatomotor (SMN) cortices and quantified simulated BOLD-like power at the stimulation frequency within the directly stimulated input networks and across indirectly engaged cortical networks (Fig. 6). We varied three gain-related parameters: cholinergic excitatory gain (α), cholinergic inhibitory gain (β), and noradrenergic filter gain (r₀).

**Figure 6.**
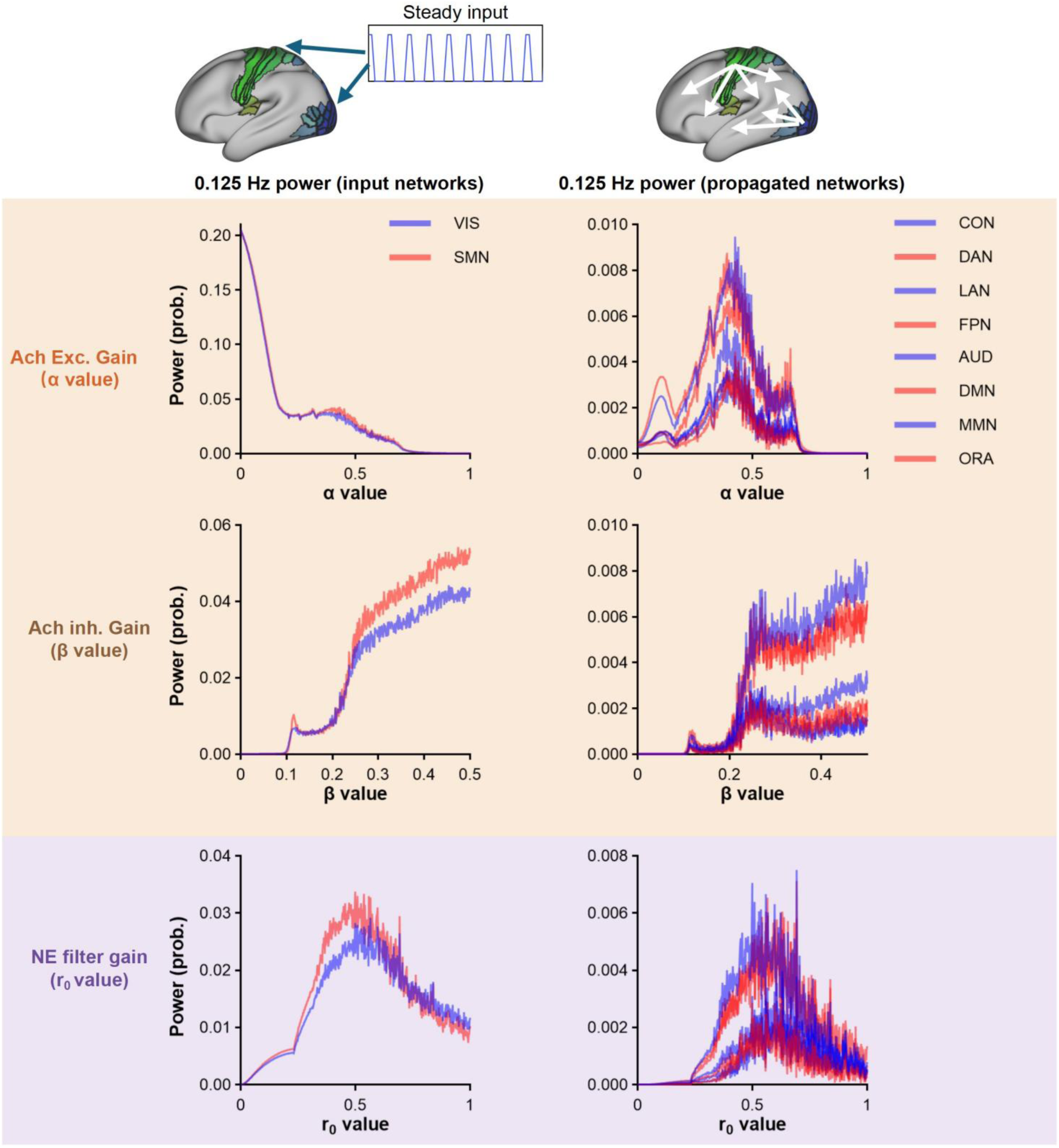
Neuromodulatory gain parameters differentially shape lfSSBR across cortical networks. Top, schematic of the whole-brain simulation. Steady-state input at 0.125 Hz was delivered to visual (VIS) and somatomotor (SMN) cortices, and activity propagated to other cortical networks through empirical structural connectivity. Bottom, simulated BOLD-like power at the stimulus-locked frequency of 0.125 Hz while varying three neuromodulatory gain parameters. Background shading denotes the neuromodulatory system being modeled: orange shading indicates cholinergic gain parameters, including excitatory gain α and inhibitory gain β; purple shading indicates the noradrenergic filter gain r₀. The left column shows power within the directly stimulated input networks (VIS and SMN), whereas the right column shows power within indirectly engaged propagated networks (CON: cingulo-opercular; DAN: dorsal attention; LAN: language; FPN: frontoparietal; AUD: auditory; DMN: default mode; MMN: multimodal; ORA: orbital-affective).

The three parameters produced distinct network profiles. Increasing α, which scales long-range excitatory input, had divergent effects depending on network class: power in directly stimulated VIS/SMN networks dropped sharply at low α values, then stabilized with a subtle local peak near α = 0.4 before declining further at higher values, whereas propagated networks exhibited a cleaner inverted-U-shaped profile also peaking near α = 0.4, indicating that intermediate excitatory gain best supports the transfer of stimulus-locked activity beyond the input networks. In contrast, increasing β, which governs inhibitory gain, enhanced 0.125-Hz power across both input and propagated networks, with a threshold-like rise at intermediate values that plateaued beyond approximately 0.25, suggesting that inhibitory tone broadly stabilizes frequency-locked responses across cortical systems. Noradrenergic filter gain r₀ showed an inverted-U-shaped profile in both network classes, with maximal stimulus-locked power at intermediate values, consistent with an optimal gain regime for temporal selectivity.

Together, these simulations show that neuromodulatory gain parameters do not simply amplify lfSSBR uniformly. Instead, cholinergic excitatory gain controlled the balance between local generation and network-wide propagation whereas cholinergic inhibitory gain strengthened frequency-locked power across networks, and noradrenergic filter gain defined an intermediate regime in which stimulus-locked power was most stable. Notably, the default parameter setting, located near the middle of the explored range, fell within this stable lfSSBR regime, suggesting that normal lfSSBR may depend on balanced rather than maximal neuromodulatory gain.

## Discussion

Within the GLM framework, task engagement is traditionally conceptualized as the superposition of transient evoked responses onto an ongoing baseline of spontaneous brain activity (Huang et al., 2017; Northoff et al., 2024). On this view, slow periodic stimulation would add power at the task-imposed frequencies locally while leaving the rest of the spectrum — in particular the brain’s dominant intrinsic low-frequency fluctuations — largely preserved, so that the spectral change would amount to the frequency-domain footprint of the response already captured by a conventional GLM. Our findings indicate that this picture is incomplete. The stimulus-locked increase did not stand in spectral isolation within the power spectrum: it was accompanied by a frequency-specific reduction concentrated within the intrinsic infra-slow range, and this redistribution persisted after the GLM-modeled response was removed. Beyond adding transient responses onto a fixed baseline, then, the task’s temporal structure transiently reorganizes ongoing infra-slow BOLD dynamics in a frequency-specific manner that conventional activation analyses do not fully capture. On this account, the stimulus-locked response is best understood not as a local addition to an otherwise independent baseline, but as a local expression of how the brain’s ongoing intrinsic dynamics are globally reorganized — consistent with the view that spontaneous activity actively shapes, rather than passively receives, stimulus-evoked responses

In the main attention dataset, this reorganization appeared as a coordinated redistribution of spectral power away from low infra-slow frequencies and toward task-imposed frequencies. Predictable rhythmic stimulation concentrated power into narrowband lfSSBR, whereas temporally irregular stimulation produced broader power engagement across task-relevant frequencies. Across independent datasets, the precise spectral expression depended on the timing of the task: when the frequency imposed by the task was separable from the brain’s dominant low-frequency range, task engagement produced a redistribution profile in the brain’s power spectrum; when the task-imposed rhythm fell within the brain’s infra-slow range, it was expressed primarily as task-frequency alignment within the brain’s ongoing infra-slow dynamics. Together, these findings indicate that task-related BOLD activity is not only spatially distributed in its activation magnitude, but also spectrally and temporally organized according to the timing demands imposed by the task.

These findings suggest that infra-slow BOLD fluctuations should not be treated as a static low-frequency background, but as frequency-resolved dynamics that can be reorganized by task demands. In the main attention dataset, task engagement reduced power in the dominant spontaneous low-frequency range while increasing power at task-imposed frequencies, indicating a redistribution of PSD rather than a global increase or decrease in BOLD power. This pattern is broadly consistent with recent proposals that different infra-slow ranges may support distinct aspects of cortical function, with lower frequencies more closely related to intrinsic organization and relatively faster infra-slow components more sensitive to sensory or task-related processing (Gong and Zuo, 2025). In this sense, task-related infra-slow BOLD dynamics resemble frequency trade-offs observed at faster electrophysiological timescales, where cognition is often accompanied by coordinated increases and decreases across frequency bands rather than isolated modulation of a single rhythm (Goldman et al., 2002; Gonçalves et al., 2006; Scheeringa et al., 2011).

The contrast between steady-state and unsteady-state stimulation further suggests that temporal predictability determines how task demands are expressed in the BOLD spectrum. Predictable rhythmic stimulation provides a stable temporal prior, allowing ongoing dynamics to align with the imposed frequency and produce lfSSBR (Lakatos et al., 2019; Schroeder and Lakatos, 2009; Van Atteveldt et al., 2015). By contrast, temporally irregular stimulation was associated with a distributed elevation of spectral power across a broader frequency range, rather than concentration into a single task-aligned frequency. This distinction is consistent with theories of temporal prediction, which propose that regular sensory structure allows the brain to anticipate when relevant events will occur and to allocate processing resources accordingly (Wang et al., 2018). In the present data, this principle was expressed at the infra-slow BOLD timescale: temporal regularity concentrated power at task-imposed frequencies, whereas temporal irregularity produced a more distributed spectral response.

An important implication of these findings is that frequency-domain organization and conventional task-evoked activation capture overlapping but non-identical aspects of task-related BOLD dynamics. lfSSBR was spatially related to GLM-derived activation: it increased in regions showing task-positive recruitment and was inversely related to regions showing task-related deactivation, suggesting that lfSSBR reflects the organization of task engagement rather than the sign of the BOLD response alone. However, this relationship was frequency-specific and incomplete. Correlations between PSD and GLM-derived activation were strongest at task-imposed frequencies and were weak or absent at non-locked frequencies, indicating that the coupling between activation and spectral power depends on the temporal structure of the task. More critically, lfSSBR persisted after removal of GLM-modeled responses, demonstrating that lfSSBR cannot be reduced to canonical task-evoked activation. This dissociation indicates that external task stimulation or neural events do not merely “activate” specific brain regions. Instead, they reorganize intrinsic infra-slow dynamics, biasing the spectral concentration of ongoing activity toward task-relevant temporal scales (He, 2011; Wang et al., 2018). Thus, GLM and PSD analyses provide complementary views of task-fMRI: the former identifies where activity increases or decreases, whereas the latter reveals how task engagement organizes BOLD dynamics across temporal scales.

Spontaneous brain activity consumes the vast majority of the brain’s energy, whereas task-evoked activation accounts for only a small fraction of total metabolic expenditure (Raichle and Mintun, 2006; Tomasi et al., 2013). This energetic imbalance highlights a central challenge for task-fMRI: cognitively relevant signals are often embedded within a much larger background of ongoing infra-slow activity. As a result, conventional amplitude-based measures may capture only part of the task-related reorganization of BOLD dynamics, limiting the sensitivity of rest-task classification, clinical state identification, and behavioral prediction (Das et al., 2021; Uddin, 2017). With machine learning becoming increasingly important in neuroscience and neural engineering, the choice of feature representation is therefore critical: improving model performance depends not only on more powerful algorithms, but also on identifying signal dimensions that more directly index task-related brain dynamics (Biswal and Uddin, 2025). Here, we propose that lfSSBR provides such a feature. By capturing frequency-specific BOLD responses imposed by stable task timing, lfSSBR reveals task-related structure within the infra-slow spectrum that is not fully captured by conventional activation measures. This is important because slow BOLD fluctuations are often treated as background variability or nuisance structure, whereas the present findings show that their frequency-specific organization can carry information about cognitive state, behavioral efficiency, and individual identity. In this sense, lfSSBR extends the logic of frequency tagging/neural entrainment to the infra-slow fMRI domain (Lakatos et al., 2008; Toffanin et al., 2009), showing that stable temporal structure can expose task-related components of BOLD dynamics that remain partly hidden in amplitude-based analyses alone.

The Jansen-Rit simulations provide a mechanistic bridge between these empirical observations and circuit-level gain control. Rather than treating lfSSBR as a passive consequence of stronger periodic input, the model suggests that frequency-specific power concentration depends on the gain regime through which external temporal structure is filtered. This interpretation is consistent with neural mass and whole-brain modeling work showing that changes in synaptic gain, excitability, and neuromodulatory control can shift large-scale brain dynamics between distinct functional regimes (David and Friston, 2003; Jansen and Rit, 1995; Li et al., 2019; Shine et al., 2018). In particular, noradrenergic gain has been proposed to regulate the responsivity and precision of cortical responses, thereby altering the balance between functional segregation and integration, whereas cholinergic modulation of local inhibitory circuits may help maintain excitation-inhibition balance while reshaping large-scale network organization (Aston-Jones and Cohen, 2005; Coronel-Oliveros et al., 2023, 2021; Servan-Schreiber et al., 1990). In our simulations, this principle was expressed in the frequency domain: excitatory gain shaped the balance between local generation and network-wide propagation of lfSSBR, inhibitory gain strengthened frequency-locked responses across cortical networks, and noradrenergic filter gain produced an intermediate regime in which temporal selectivity was maximal. Thus, task-imposed infra-slow organization appears to arise not from maximal amplification of sensory drive, but from a balanced gain state that allows ongoing cortical dynamics to align with predictable external timing while remaining dynamically stable.

### Limitations

Several limitations should be considered. First, although the present findings indicate that task-imposed spectral features are robust across datasets and persist after removal of GLM-modeled responses, BOLD power spectra remain an indirect measure of neural activity. The observed stimulus-locked BOLD peaks should not be interpreted as direct neural oscillations, but as frequency-specific organization of hemodynamically filtered population activity. We note, however, that in our earlier work, frequency-specific effects persisted after HRF deconvolution, suggesting that they are not solely produced by hemodynamic filtering and likely reflect a neuronal contribution (Wang et al., 2016, 2015, 2014). Although analyses of raw PSD, GLM residuals, and framewise displacement argue against explanations based solely on normalization, canonical activation, or head motion, other physiological sources such as respiration, cardiac activity, vascular reactivity, and individual differences in hemodynamic filtering may still contribute to infra-slow BOLD spectra. Second, the three datasets differed in task modality, temporal structure, stimulation frequency, and sample characteristics. This heterogeneity strengthens the general claim that task timing can organize BOLD spectral power, but it also limits direct quantitative comparison across datasets and prevents us from concluding that different task structures produce quantitatively identical redistribution profiles. Third, the GLM-residual analysis shows that stimulus-locked peaks are not fully reducible to canonical task-evoked activation, but the analysis of the residuals cannot completely separate frequency-specific temporal organization from all possible task-related hemodynamic components, especially when the GLM regressors and spectral peaks share the same temporal structure. Finally, the Jansen-Rit simulations provide a sufficiency account rather than a direct parameter inversion of empirical data. The gain parameters were used to test whether changes in excitability, inhibition, and filtering can generate the observed frequency-specific organization, but they should not be taken as direct evidence for measured cholinergic or noradrenergic activity in participants. Future work combining task-frequency fMRI with physiological monitoring, pharmacological manipulation, pupillometry, EEG/MEG, or individualized model fitting will be needed to establish the neural and neuromodulatory sources of spectral features more directly.

## Conclusion

In conclusion, the present study shows that task engagement with temporally structured events reorganizes infra-slow BOLD dynamics in a frequency-specific manner. Across independent datasets, task timing shaped the redistribution within the BOLD power spectrum’s whole frequency range, producing either lfSSBR or broader redistribution across task-relevant frequencies. lfSSBR peaks were not fully reducible to conventional GLM-derived activation, and carried information about cognitive state, behavioral performance, and individual identity. Mechanistically, Jansen-Rit simulations suggest that such frequency-specific organization can arise from balanced changes in excitatory, inhibitory, and neuromodulatory filter gain, rather than from simple amplitude amplification by the external input. Together, these findings suggest that task-based fMRI can reveal not only the localization of specific cognitive functions in particular regions, but also how the task’s temporal structure transiently reorganizes the BOLD dynamics’ power spectrum in a task- and frequency-specific way.

## Methods

### Subjects

#### UESTC Sample

Fifty healthy adults (31 males, 19 females; mean age ± SD = 23.12 ± 3.90 years; range: 18-45 years) were recruited as the primary sample in the University of Electronic Science and Technology of China (UESTC). All participants were right-handed, had normal or corrected-to-normal vision, and reported no history of neurological, psychiatric, or major medical conditions. Three participants were excluded due to high task error rates (>30%) or unusually long reaction times (>3 SD above the group mean), resulting in a final sample of 47 individuals. Written informed consent was obtained from each participant before the experiment. The study was approved by the Research Ethics Committee of the School of Life Science and Technology at UESTC and was conducted in accordance with the Declaration of Helsinki.

#### HZNU Sample

We used an independent dataset of 25 right-handed adults (13 males, 12 females; age: 20-29 years) previously reported in (Huang et al., 2017). Participants had no history of neurological or psychiatric disorders, as confirmed through standard MRI safety screening. Written informed consent was obtained from each subject prior to the experiment. The study was approved by the Ethics Committee of the Center for Cognition and Brain Disorders (CCBD), Hangzhou Normal University.

#### ZJU Sample

The second replication dataset included 33 adults (17 males, 16 females; age range: 21-36 years) who completed a task-based fMRI experiment previously reported in (Liu et al., 2024). All participants were right-handed and had normal or corrected-to-normal vision. All procedures were approved by the Ethics Review Committee of Zhejiang University and were conducted in accordance with the Declaration of Helsinki. Written informed consent was obtained from all participants before the start of the study, and participants were compensated for their time. Four participants were excluded from the final analysis: three due to excessive head motion (mean framewise displacement > 0.35 mm) and one due to missing imaging data, resulting in a final sample of 29 individuals.

### Task procedure

#### UESTC dataset

Two visual detection tasks [regularly presented (steady-state task) vs. unregularly presented (unsteady-state task)] lasting for 10 minutes and an equal-length resting scan were counterbalanced between subjects. During the resting scan, participants were required to remain motionless, focus their eyes on a white crosshair against black background, stay awake, and not think of anything in particular. During task presentations, participants were asked to focus on a black crosshair located at the center of screen during the entire task. They were required to press the left key with the left thumb as accurately and fast as possible if a yellow disc appeared at the left of the crosshair, or to press the right key with the right thumb if a yellow disc appeared at the right of the crosshair. The diameter of disc is 1 cm with the horizontal view angle of 13.5° from the crosshair. As shown in Fig. 1, in each stimuli series, the black crosshair always stayed at the center of screen whereas the disc was presented on the gray background for 0.1 s. At the steady-state task, the left disc was presented every 12 s while the right one was presented every 8 s, forming two visual flows of 0.083 Hz and 0.125 Hz. At the unsteady-state task, the left disc was presented with a mean pace of 12 s (trial length ranges from 8 to 16 s) while the right one was presented with a mean pace of 8 s (trial length ranges from 4 to 12 s), forming a range of visual flows between 0.0625 Hz to 0.25 Hz. Because both visual and motor processes have contralateral advantage, this design could test whether spectral feature is task-related by detecting greater responses in contralateral cortices. The procedure was performed with E-Prime 2.0 software (http://www.pstnet.com; Psychology Software Tools).

#### HZNU dataset

This scanning included a 6-minute resting run and a 20-minute task run originally reported in Huang et al. (2017). Before scanning, participants completed a 360-item questionnaire assessing autobiographical and factual knowledge, from which 120 individualized questions were selected (Huang et al., 2014). During the task, 20 auditory questions were presented with very long inter-trial intervals (52-60 s), allowing responses to return fully to baseline and minimizing hemodynamic overlap. Participants indicated “yes”/“no” responses via button press.

#### ZJU dataset

The dataset employed a high-resolution visual motion detection task. Each run comprised 13 blocks (20 s each), consisting of 10 s baseline followed by 10 s drifting-grating stimulation. Stimuli were sinusoidal luminance-modulated gratings (50% contrast; 1 cycle/° spatial frequency; 4°/s motion speed) moving either leftward or rightward and presented at two diameters (2° and 10°), yielding four stimulus conditions. Gratings were displayed centrally on a gray background (56 cd/m²) with blurred edges (raised cosine, 0.3°). Stimulus onset and offset were temporally smoothed using a Gaussian envelope, with duration defined as 1 SD of the Gaussian and adaptively adjusted via an interleaved three-down/one-up staircase procedure. Duration thresholds for each stimulus size were estimated from a 160-trial block by fitting a cumulative Gaussian psychometric function and extracting the 75% correct point. Practice trials preceded the experiment, and auditory feedback was provided for incorrect responses.

### Imaging data acquisition

#### UESTC dataset

MRI data were acquired on a 3T GE 750 scanner equipped with high-speed gradients, using an 8-channel prototype quadrature birdcage head coil. Foam padding was used to minimize head motion. Functional images were acquired using a gradient-echo echo-planar imaging (EPI) sequence with the following parameters: repetition time/echo time (TR/TE) = 2000/30 ms, flip angle = 90°, bandwidth = 250 Hz/pixel, 43 axial slices, slice thickness = 3.2 mm with no gap, matrix = 64 × 64, field of view (FOV) = 220 mm, and 300 volumes.

#### HZNU dataset

MRI data were acquired on a 3T GE Discovery MR750 scanner using an 8-channel head coil. Whole-brain functional images were acquired using a gradient-echo EPI sequence with the following parameters: TR/TE = 1000/25 ms, flip angle = 76°, 21 slices, slice thickness = 6 mm with no gap, matrix = 64 × 64, and FOV = 210 mm. The resting-state run comprised 360 volumes, corresponding to 6 min, and the task run comprised 1184 volumes, corresponding to 19 min 44 s.

#### ZJU dataset

MRI data were acquired on a 7T Siemens whole-body scanner using a 32-channel Nova Medical head coil. Functional images were acquired using an EPI sequence with 1.5-mm isotropic resolution in transverse orientation and the following parameters: TR/TE = 2000/20.6 ms, flip angle = 70°, 90 slices, and 160 volumes. Participants were instructed to keep their eyes closed. High-resolution structural images were acquired using an MP2RAGE sequence with the following parameters: TR/TI1/TI2 = 5000/901/3200 ms and 0.7-mm isotropic resolution.

### Imaging data preprocessing

UESTC and HZNU dataset: Functional images were preprocessed using *fMRIPrep* (version 23.0.2) with default parameters (Esteban et al., 2019). Preprocessed outputs were generated in the fsaverage5 surface space and stored as CIFTI files for subsequent analyses. The first four volumes of each run were discarded to allow for signal stabilization. Nuisance regression was then performed to remove confounding effects associated with six rigid-body head motion parameters, including three translations and three rotations, their first derivatives, framewise displacement (FD), and mean signals from white matter, cerebrospinal fluid, and the whole-brain global signal. All participants met the head motion criterion of mean FD < 0.35 mm. No temporal band-pass filtering was applied, as the present study focuses on the full infra-slow power spectrum of the BOLD signal and its redistribution across frequencies from rest to task. Cortical time series were extracted using the Glasser et al. (2016) multimodal parcellation. For each of the 360 regions, vertex-wise signals were averaged to yield a single representative regional time series (Glasser et al., 2016).

#### ZJU dataset

Preprocessing of the ZJU dataset was conducted using the Data Processing Assistant for Resting-State fMRI (DPARSF) (Yan et al., 2016). The first four volumes were removed, followed by slice-timing correction and head-motion realignment. Functional images were normalized to the standard EPI template and resampled to 3 × 3 × 3 mm³ isotropic resolution. Nuisance regression included linear trends, white matter signals, cerebrospinal fluid signals, the whole-brain global signal, and Friston’s 24 head-motion parameters. Three participants were excluded due to excessive head motion (mean FD > 0.35 mm). Finally, gray matter was parcellated into 246 regions using the Human Brainnetome Atlas, and the mean time series within each region was extracted to represent regional activity.

Because the present study quantified frequency-specific BOLD power across cortical regions, spatially diffuse fluctuations could contribute to apparent spectral effects, particularly when assessing task-frequency responses across the cortex. We therefore applied global signal regression consistently across all three datasets to reduce variance shared broadly across the brain, including components related to systemic physiology, residual motion, and other non-specific global fluctuations (Fox et al., 2005; Murphy and Fox, 2017).

### Spectral power analysis

The power spectral density (PSD) of the time series from each of the 360 brain regions was computed using the *periodogram* function in MATLAB, with FFT length set to the next power of two greater than or equal to the scan length. Frequency bins from 0.01 Hz up to the Nyquist frequency were retained for subsequent analyses. Each region’s PSD was expressed as relative power by dividing the power at each frequency bin by the total power summed across the retained frequency range.

Stimulus-locked spectral power (lfSSBR) was quantified around the two prominent power peaks observed near the fundamental stimulation frequencies in the main UESTC dataset, corresponding approximately to 0.083 Hz and 0.125 Hz. For each frequency, power was averaged across the nearest frequency bin and its two adjacent bins to reduce the influence of spectral leakage and small mismatches between the stimulation frequency and the discrete FFT grid. This three-bin narrowband estimate was then used as the stimulus-locked spectral feature in subsequent analyses.

For multivariate analyses, including SVM classification, elastic net regression, and individual identification, relative-power spectra were z-scored within each region and scan across the retained frequency bins. This step rescaled the relative-power values to a numerically stable range for model fitting, because relative-power features are bounded and small and may otherwise affect scale-sensitive models such as SVM and elastic net regression.

For all subsequent analyses, each of four frequency features of interest was operationalized as a single mean-power value per region per scan: the mean across the three bins of each narrowband window centered on 0.083 Hz and 0.125 Hz, and the mean across all bins within the low infra-slow range of 0.01-0.04 Hz and the broad task-relevant range of 0.0625-0.25 Hz, respectively. For each brain region, as well as for the whole-brain averaged PSD, condition differences in mean power at each frequency feature were assessed using paired t-tests, with multiple comparisons across conditions, frequency features, and brain regions controlled using false discovery rate correction at *q* < 0.05. The same procedure was applied to the HZNU and ZJU replication datasets.

### Frequency-resolved redistribution analyses

Without predefining frequency bands, we performed three complementary frequency-resolved analyses on the whole-brain averaged relative-power spectra, separately for the steady- and unsteady-state conditions of the UESTC dataset. (1) To localize task-related power changes, we contrasted task and rest at each frequency bin with a paired t-test (positive t indicating task > rest). (2) To test whether stimulus-locked power traded off against specific frequencies across individuals, we correlated, across subjects, each subject’ssubject’s lfSSBR power (whole-brain power summed across the 0.083 Hz and 0.125 Hz bins) with whole-brain power at every other bin (Pearson); the effect is displayed as the regression slope (β), with negative β indicating lower power at a given frequency in subjects with higher lfSSBR power. (3) To capture the same coupling within individuals, we segmented each subject’ssubject’s BOLD time series into 120 s windows (60 volumes; 4 s / 2-volume step), recomputed the whole-brain relative-power spectrum per window, and regressed window-wise power at every bin on the window-wise lfSSBR time course (yielding within-subject β and Pearson r); group inference used a one-sample t-test of the Fisher z-transformed correlations against zero.

In all three analyses, multiple comparisons across frequency bins were controlled using one-dimensional cluster-based permutation testing (Cox et al., 2020; Maris and Oostenveld, 2007; Nichols and Holmes, 2002) (10,000 permutations; cluster-forming threshold p < 0.05, two-tailed; cluster-level threshold p < 0.1), with cluster mass defined as the summed bin-wise statistic within each contiguous suprathreshold cluster and the null distribution built from the maximum cluster mass across permutations. The null distribution was generated by sign-flipping the within-subject task-rest difference (analysis 1), permuting lfSSBR values across subjects (analysis 2), and sign-flipping the Fisher-z values across observations (analysis 3).

### The relationship between spectral features and task-evoked activation

Task-evoked activation was estimated using a general linear model (GLM) fitted separately to each task condition. For each condition (steady-state and unsteady-state), two stimulus-onset regressors were constructed — one for the left and one for the right visual stimulus stream — using the actual stimulus onset times of each stream within that condition. In the steady-state condition, onsets occurred periodically at 0.083 Hz and 0.125 Hz; in the unsteady-state condition, the identical modeling procedure was applied to each stream’s aperiodic onset sequence. Stimulus onsets were modeled as impulse functions and convolved with the canonical hemodynamic response function using the *spm_hrf* function from the SPM toolbox. A constant term was included as a nuisance regressor. Each condition-specific design matrix was fitted to the corresponding segment of the regional BOLD time series, yielding left-stream and right-stream beta estimates for each brain region.

The statistical significance of task-evoked activation was assessed using one-sample t-tests on regional GLM beta estimates across subjects, with multiple comparisons across the 360 regions controlled using false discovery rate correction at *q* < 0.05. To examine the relationship between spectral features and conventional task-evoked activation, we performed across-subjects Pearson correlations on a per-region basis. For each brain region, the vector of GLM beta estimates across subjects was correlated with the corresponding vector of regional relative-power values across the same subjects, yielding one Pearson correlation coefficient per region. Correlations were computed separately for the left-stream and right-stream beta estimates, separately for the steady-state and unsteady-state conditions, and separately for each PSD feature of interest (0.01-0.04 Hz, 0.0625-0.25 Hz, 0.083 Hz, and 0.125 Hz, as defined in the spectral analysis). Multiple comparisons across brain regions were controlled using false discovery rate correction at *q* < 0.05.

To test whether stimulus-locked BOLD peak effects could be reduced to canonical task-evoked responses, the two stimulus-onset regressors were regressed out from the task-state BOLD signals, and the same spectral analyses were repeated on the residual time series. As a control analysis, the same task design matrices were also fitted to resting-state BOLD signals from the same subjects, for which no true task-evoked response was expected. Beta estimates from these resting-state control GLMs were not interpreted as activation; instead, this analysis tested whether the residual spectral feature patterns could arise from spurious fitting of task regressors to ongoing BOLD fluctuations.

### Support vector machine (SVM) analysis

To classify brain states based on spectral features, we applied support vector machine (SVM) pattern analysis to distinguish rest, steady-state task, and unsteady-state task conditions. Classification was performed separately for four spectral features. For each feature set, the input vector for each sample consisted of spectral power values across all brain regions, with each subject contributing one sample per condition.

A multiclass error-correcting output codes model with one-vs-one binary SVM learners was implemented using MATLAB’s fitcecoc function. Model performance was evaluated using leave-one-subject-out cross-validation, in which all condition samples from one subject were held out for testing and the model was trained on the remaining subjects. In each fold, the trained ECOC-SVM model predicted the condition labels of the held-out subject’s samples. Performance was summarized using overall classification accuracy and class-wise area under the receiver operating characteristic curve (AUC). Class-wise AUC was computed in a one-vs-rest manner using the classification scores returned by the model and MATLAB’s perfcurve function.

Pairwise differences in AUC across spectral features were tested using DeLong’s test, and p values were corrected for multiple comparisons across all feature pairs and classes using the false discovery rate procedure, with significance defined as *q* < 0.05.

### Elastic net regression for behavioral prediction

Elastic net regression was used to test whether spectral power redistribution predicted individual mean reaction time (RT) across all trials. Paralleling the SVM analysis, models were trained separately for each spectral feature set using 360 regional PSD features. Elastic net regularization was chosen because the feature space contained many spatially correlated regional predictors relative to the sample size; its L1 component promotes sparse feature weighting, whereas its L2 component stabilizes coefficient estimates under multicollinearity (Zou and Hastie, 2005).

Model fitting was implemented in MATLAB using the *lasso* function, which minimizes the following objective with respect to the regression coefficients β:

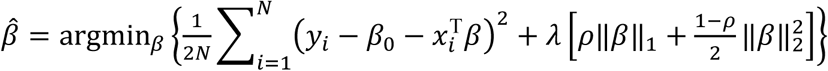

where *N* is the number of training subjects, *y_*i*_* is the observed mean RT of subject *i*, **x_*i*_** ∈ ℝ³⁶⁰ is the corresponding regional feature vector, β₀ is the intercept, λ ≥ 0 controls the overall regularization strength, and ρ ∈ (0, 1] controls the L1/L2 mixing ratio. Here, ρ is used in place of MATLAB’s α notation to avoid confusion with the neuromodulatory gain parameter α which will be introduced later. ρ = 1 corresponds to pure Lasso regularization, whereas smaller ρ values introduce stronger Ridge regularization. The mixing parameter ρ was searched over 0.05-1.00 in steps of 0.05. For each ρ, λ was searched over the default 100-value sequence automatically generated by *lasso* function.

Predictive performance was evaluated using nested leave-one-out cross-validation. In each outer fold, one subject was held out as the test sample and the remaining subjects formed the training set. Within the training set, features were standardized using the training-set mean and standard deviation, and the same parameters were applied to the held-out subject to prevent information leakage. Hyperparameters were selected within the outer training set using repeated 10-fold cross-validation, repeated 10 times with different fold partitions. The ρ and λ pair with the lowest average inner-CV mean squared error was then used to refit the model on the full outer training set and predict RT for the held-out subject.

Predictions were aggregated across all outer folds. Model performance was quantified using the cross-validated coefficient of determination, *Q*^2^ = 1 − SSE/SST, where *Q*^2^ > 0 indicates better-than-mean prediction. Statistical significance for each *Q*^2^value was assessed using permutation testing, in which RT labels were randomly shuffled 10000 times and the full nested-CV procedure was repeated to generate an empirical null distribution.

Pairwise differences in predictive performance across feature sets (frequencies, PSD vs. GLM) were assessed using paired sign-flip permutation tests on subject-level squared prediction errors. For each pair of feature sets, squared errors (SE) were computed per subject, and the observed test statistic was the mean paired difference of mean SE. To generate the null distribution, the sign of each subject’s paired difference was independently flipped with probability 0.5 and the mean recomputed; this was repeated 10,000 times. Two-sided p-values were computed. 95% confidence intervals for the MSE of each feature set and for mean paired MSE differences were obtained via paired bootstrap resampling of subject indices (10,000 iterations). P-values across the pairwise feature comparisons were corrected for multiple comparisons using false discovery rate at q < 0.05.

### Individual identification based on PSD spatial patterns

To assess whether spectral power redistribution carried subject-specific information, we performed an individual identification analysis based on regional PSD patterns across 6 task runs in HZNU dataset, which was the only dataset with multiple task runs per subject suitable for between-run identification, following the method suggested by (Finn et al., 2015). For each frequency feature, each subject was represented by a 360-dimensional PSD pattern across cortical regions. Identification was performed for each ordered pair of runs, with one run used as the reference set and the other as the query set. For each query subject, spatial Pearson’s correlations were computed between their PSD pattern and the PSD patterns of all subjects in the reference run. The predicted identity was assigned to the reference subject with the highest spatial correlation. Identification was considered correct when the predicted identity matched the true identity of the query subject. Subjects with missing values in either run were excluded from the corresponding run-pair comparison. Identification accuracy was calculated as the proportion of correctly identified subjects for each run pair and frequency feature, and then averaged across all off-diagonal run pairs.

### Activation-based prediction models

To compare stimulus-locked BOLD peaks-related power redistribution with conventional task-evoked activation, we repeated the applicable multivariate analyses using GLM-derived activation features. For each subject and task condition, beta estimates from the two stimulus-onset regressors, corresponding to the 0.083-Hz and 0.125-Hz stimulus streams, were extracted across all 360 brain regions to form activation-based feature vectors. Because GLM beta estimates are defined relative to a task design and have no equivalent measure in the resting-state condition, activation features were not used for the three-class SVM state-classification analysis.

Activation-based features were instead submitted to the same elastic net regression pipeline used for PSD features to predict individual reaction time. In addition, we repeated the individual identification analysis using spatial patterns of GLM beta estimates across the 360 regions, following the same reference-query run-matching procedure described above. Differences in behavioral prediction performance between PSD-based and GLM-based models were assessed using a permutation-based test for dependent prediction correlations.

### Computational modeling: Jansen-Rit neural mass network

The Jansen-Rit framework (Jansen and Rit, 1995) models a cortical area as a local neural mass circuit comprising three interacting neural populations: pyramidal neurons, excitatory interneurons, and inhibitory interneurons. Within each cortical area, pyramidal cells receive synaptic input from both excitatory and inhibitory interneuron populations and, in turn, project back to them, forming a canonical local feedback loop. This local architecture captures the recurrent excitation and inhibition that shape population-level cortical dynamics.

To model large-scale brain dynamics, we extended the local Jansen-Rit circuit to a network of *N* = 360 coupled cortical regions. Long-range cortico-cortical projections were weighted by a publicly available group-average structural connectivity matrix derived from probabilistic diffusion tractography on 1065 Human Connectome Project subjects (Rosen and Halgren, 2021). The connectivity matrix *W* was then used to couple regional neural masses through a long-range pyramidal projection channel between cortical regions. In this network formulation, pyramidal output from one region propagates through the structural connectome and provides excitatory drive to the pyramidal population of downstream regions.

Following Coronel-Oliveros et al. (2021), we incorporated two modifications relative to the original Jansen-Rit formulation to make the model suitable for large-scale cortico-cortical interactions. First, an additional pyramidal projection state, *x*_3_, was introduced to represent long-range cortico-cortical output. This state is driven by the same pyramidal firing-rate signal as the local pyramidal channel but is integrated with a slower effective timescale, implemented here as *a*_*d*_ = 0.5*a*. This additional filtering captures the delayed and smoothed influence of long-range excitatory projections, consistent with the idea that distal cortico-cortical inputs are integrated through dendritic filtering mechanisms. Second, a local inhibitory-to-excitatory interneuron connection was added to the canonical Jansen-Rit motif. This short-range inhibitory feedback provides a stabilizing mechanism that constrains recurrent excitation, helps preserve local excitation-inhibition balance, and allows the network to transition between more segregated and more integrated dynamical regimes.

Each node *i* was therefore described by four post-synaptic potential-like state variables: the pyramidal channel 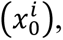 excitatory-interneuron channel 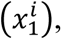 inhibitory-interneuron channel 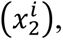 and long-range coupling channel 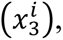 together with their first-order temporal derivatives. A sigmoid transfer function *S*(⋅) converted mean membrane potential into mean firing rate. Each region additionally received a tonic background input *p*, perturbed by additive Gaussian noise with amplitude *σ*. External stimulation *u*_*i*_(*t*) was introduced as an additive drive to the background input. The model equations are written below:

#### Sigmoid transfer function

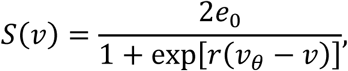

where *e*_0_ is the half-maximum firing rate, *v*_*θ*_ is the half-activation membrane potential, and *r* controls the slope of the sigmoid function.

#### Long-range (cortico-cortical) input

After removing self-connections, the structural connectivity matrix was normalized as

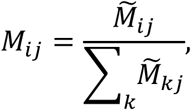

where *M̃* denotes the empirical structural connectivity matrix after setting diagonal entries to zero. The long-range input to node *i* was then given by

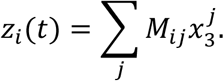

#### External input

The tonic input to each node was defined as

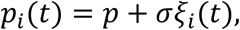

where *ξ*_*i*_(*t*)is Gaussian noise.

#### Pyramidal firing rate is obtained via the sigmoid transfer function as

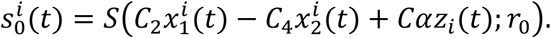

#### Pyramidal channel (*x*_0_)

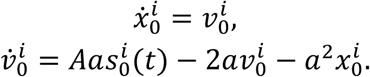

This channel represents the local pyramidal population driven by the combined effects of excitatory interneuron input, inhibitory interneuron input, and long-range cortico-cortical input.

#### Excitatory-interneuron channel (*x*_1_)

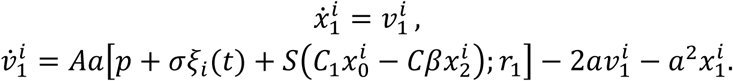

This channel represents the excitatory interneuron population. The term 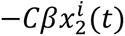 implements local inhibitory feedback onto the excitatory-interneuron pathway.

#### Inhibitory-interneuron channel (*x*_2_)

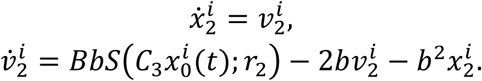

This channel represents the inhibitory interneuron population driven by pyramidal activity.

#### Long-range coupling channel (*x*_3_)

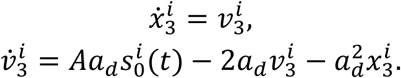

This channel carries pyramidal output into the large-scale structural network and mediates cortico-cortical propagation.

In the Jansen-Rit model, the maximum firing rate was set to *e*_0_ = 2.5 Hz, the half-activation membrane potential to *v*_*θ*_ = 6 mV, and the default sigmoid slopes to *r*_0_ = *r*_1_ = *r*_2_ = 0.56 mV^−1^. Excitatory and inhibitory synaptic gains were set to *A* = 3.25 mV and *B* = 22 mV, respectively. The corresponding inverse time constants were *a* = 100 s^−1^ for excitatory synapses and *b* = 50 s^−1^ for inhibitory synapses. The long-range coupling channel used a separate inverse time constant *a*_*d*_ = 50 s^−1^, corresponding to *a*_*d*_ = 0.5*a*. The tonic background input was set to *p* = 2, and the amplitude of Gaussian noise was set to *σ* = 2. Local intra-column connection strengths were defined relative to a base coupling constant *C* = 135, following the standard Jansen-Rit ratios: *C*_1_ = *C*, *C*_2_ = 0.8*C*, *C*_3_ = 0.25*C*, *C*_4_ = 0.25*C*.

Here, *C*_1_ and *C*_3_ scale projections from the pyramidal population to excitatory and inhibitory interneurons, respectively, whereas *C*_2_ and *C*_4_ scale feedback projections from excitatory and inhibitory interneurons back to the pyramidal population.

Three gain-related parameters were incorporated to model neuromodulatory influences on large-scale dynamics. The cholinergic excitatory gain *α* scaled the effective strength of long-range excitatory input to the pyramidal population through the term *Cαz*_*i*_(*t*). Larger values of *α* therefore increase the sensitivity of each region to inter-regional input and promote the propagation of stimulus-driven activity across the structural network. The cholinergic inhibitory gain *β* scaled inhibitory feedback onto the excitatory-interneuron channel through the term 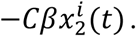 This parameter regulates how strongly local inhibition suppresses recurrent excitatory drive, thereby modulating local excitation-inhibition balance. Finally, following Coronel-Oliveros et al. (2021), we treated *r*_0_ as a phenomenological proxy for noradrenergic modulation of pyramidal response gain. By changing the slope of the pyramidal sigmoid transfer function, *r*_0_ modulates how sharply membrane-potential input is transformed into firing-rate output and determines how sensitively the pyramidal population responds to local and long-range input.

Together, *α*, *β*, and *r*_0_ provided a mechanistic axis for testing how neuromodulatory control of long-range excitation, local inhibitory stabilization, and pyramidal response gain shapes stimulus-locked spectral redistribution across the structural connectome. In parameter-sweep analyses, these gain-related parameters were varied systematically while other model parameters were held fixed at their default values.

The neural mass model was integrated using a forward Euler scheme with a time step of *dt* = 10^−3^ s. Each simulation included an initial equilibration period of 60 s which we discard, followed by 600 s of simulated neural activity used for subsequent analyses. The pyramidal firing-rate output 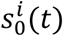 from each region was stored after temporal downsampling, yielding an effective sampling interval of 0.01 s. This pyramidal output was then used as the neural drive to a Balloon-Windkessel hemodynamic model to generate BOLD-like signals.

### Balloon-Windkessel model

For each region, the neural drive *n*_*i*_(*t*) was used as input to a four-state hemodynamic system comprising the vasodilatory signal *s*_*i*_(*t*), blood inflow *f*_*i*_(*t*), venous blood volume *v*_*i*_(*t*), and deoxyhemoglobin content *q*_*i*_(*t*). The hemodynamic equations were integrated using the same forward Euler scheme:

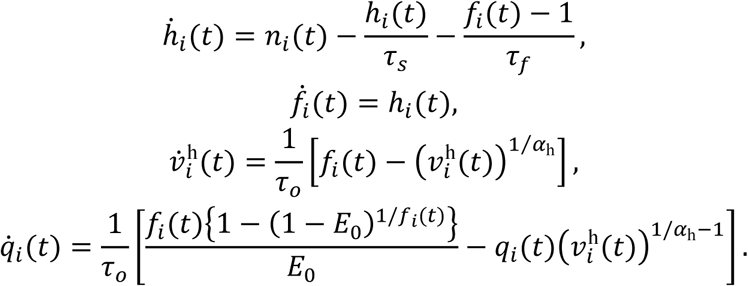

Here, *τ*_*s*_ is the signal decay constant, *τ*_*f*_ controls autoregulatory feedback from blood inflow, and *τ*_*o*_ is the mean transit time governing venous blood volume and deoxyhemoglobin dynamics. The parameter *α*_ℎ_ controls vessel stiffness, and *E*_0_ denotes the resting oxygen extraction fraction. Initial conditions were set to *s*_*i*_(0) = 0.1, *f*_*i*_(0) = 1, *v*_*i*_(0) = 1, and *q*_*i*_(0) = 1 for all regions.

The BOLD signal was computed from the simulated hemodynamic states as

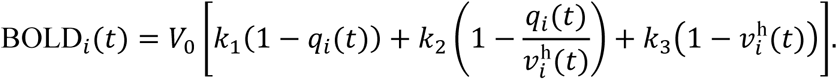

where *V*_0_ is the resting venous blood volume fraction. The coefficients were defined as

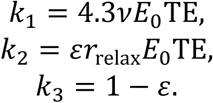

The hemodynamic parameters were set to *τ*_*s*_ = 0.65 s, *τ*_*f*_ = 0.41 s, *τ*_*o*_ = 0.98 s, *α*_ℎ_ = 0.32, *E*_0_ = 0.4, TE = 0.04 s, and *V*_0_ = 0.04. The frequency offset was set to *ν* = 40.3 s^−1^, the intravascular relaxation parameter to *r*_*relax*_ = 25 s^−1^, and the intra- to extravascular signal ratio to *ε* = 0.5. The first 60 s of the simulated BOLD-like signal were discarded to remove transient hemodynamic effects before spectral analyses.

### Parameter sweep and robustness analysis

Two complementary parameter-sweep analyses were performed using the JR + Balloon-Windkessel model. The first analysis tested the robustness of the stimulus-locked BOLD peaks phenomenon to perturbations of canonical JR biophysical parameters, whereas the second examined how three neuromodulatory gain parameters shape spectral power redistribution. All sweep simulations used the same stimulation protocol as in the empirical paradigm. External stimulation was implemented as 1-s rectangular pulses (amplitude = 0.8) added to the background drive *p*_*i*_(*t*). In the steady-state condition, pulses were delivered periodically at 0.125 Hz. In the unsteady-state condition, pulses were delivered as an aperiodic sequence with inter-pulse intervals randomly drawn from a uniform distribution between 4 and 16 s, corresponding to an approximate broadband temporal profile of 0.0625-0.25 Hz.

#### Robustness sweep

To test whether the simulated spectral feature phenomenon depended on the precise canonical values of the JR model parameters, eight biophysical parameters, including *a, b, A, B, C1, C2, C3, C4,* were swept independently at the single-node level over a ±20% range around their canonical values, with 501 evenly spaced values per parameter. Neuromodulatory parameters were held fixed at their reference values (*α* = 0.5, *β* = 0.25, *r*_0_ = 0.56 mV^−1^) throughout the robustness sweep. This analysis was designed to assess whether stimulus-frequency-locked power redistribution remained stable under moderate perturbations of the canonical JR parameter regime.

#### Neuromodulator sweep

Three one-dimensional parameter sweeps were then performed by varying one neuromodulatory parameter at a time while holding the other two fixed at intermediate reference values (*α* = 0.5, *β* = 0.25, *r*_0_ = 0.56 mV^−1^), following the integration regime described by Coronel-Oliveros et al. (2021). The swept parameters were cholinergic excitatory gain *α* ∈ [0,1], cholinergic inhibitory gain *β* ∈ [0,0.5], and noradrenergic response-gain parameter *r*_0_ ∈ [0,1] mV^−1^. Each sweep comprised 200 evenly spaced parameter values. To assess how long-range structural coupling shapes neuromodulatory effects, each sweep was performed under two configurations: a single-node model without inter-regional coupling, and a whole-brain network model in which all 360 cortical regions were coupled through the column-normalized structural connectivity matrix *M̃*.

In the whole-brain configuration, external stimulation was injected only into ROIs belonging to the visual (VIS) and somatomotor (SMN) networks. This choice reflected the empirical paradigm, in which visual stimulation engages sensory processing and motor preparation for button-press responses. To characterize how stimulus-evoked power redistribution propagated from directly stimulated regions to the rest of the cortex, the 360 Glasser’s ROIs were assigned to functional networks following the Cole-Anticevic Brain-wide Network Partition (Ji et al., 2019). Networks were then grouped into two categories: input networks, comprising VIS and SMN, which received direct external stimulation; and propagated networks, comprising the cingulo-opercular (CON), dorsal attention (DAN), language (LAN), frontoparietal (FPN), auditory (AUD), default mode (DMN), multimodal (MMN), and orbital-affective (ORA) networks, in which power redistribution arose solely through long-range structural coupling. For each neuromodulator sweep, relative power at the stimulus fundamental frequency of 0.125 Hz was averaged across all ROIs within each network and expressed as a function of the swept parameter, separately for input and propagated networks. This decomposition dissociated the direct effect of stimulation at sensory/motor sites from network-mediated propagation effects in non-stimulated regions.

#### PSD computation and statistical analysis

For each swept parameter value and each stimulation condition, simulated BOLD-like signals were generated, downsampled to *TR* = 2 s, matching the empirical fMRI sampling rate, and submitted to power spectral density estimation using MATLAB’s periodogram function with an FFT length of 512. PSD values were expressed as relative power by normalizing each spectrum to sum to one across the retained frequency range from 0.01 Hz to the Nyquist frequency, consistent with the empirical analysis. The relationship between each swept parameter and simulated relative power was quantified frequency-by-frequency using distance correlation, which captures both linear and nonlinear dependencies. This was particularly relevant because neuromodulatory parameters can exhibit nonlinear relationships with network-level dynamics. Resulting *p*-values were corrected across frequency bands using the FDR procedure with *q* < 0.05.

## References

Abram, S.V., Helwig, N.E., Moodie, C.A., DeYoung, C.G., MacDonald, A.W., Waller, N.G., 2016. Bootstrap Enhanced Penalized Regression for Variable Selection with Neuroimaging Data. Front. Neurosci. 10. 10.3389/fnins.2016.00344

Ao, Y., Klar, P., Catal, Y., Wang, Y., Northoff, G., 2025. Infra-slow scale-free dynamics modulate the connection of neural and behavioral variability during attention. Commun. Biol. 8, 1057.

Aston-Jones, G., Cohen, J.D., 2005. AN INTEGRATIVE THEORY OF LOCUS COERULEUS-NOREPINEPHRINE FUNCTION: Adaptive Gain and Optimal Performance. Annu. Rev. Neurosci. 28, 403–450. 10.1146/annurev.neuro.28.061604.135709

Bianciardi, M., Fukunaga, M., Van Gelderen, P., Horovitz, S.G., De Zwart, J.A., Duyn, J.H., 2009. Modulation of spontaneous fMRI activity in human visual cortex by behavioral state. NeuroImage 45, 160–168. 10.1016/j.neuroimage.2008.10.034

Biswal, B.B., Uddin, L.Q., 2025. The history and future of resting-state functional magnetic resonance imaging. Nature 641, 1121–1131. 10.1038/s41586-025-08953-9

Bola, M., Sabel, B.A., 2015. Dynamic reorganization of brain functional networks during cognition. Neuroimage 114, 398–413.

Buzsáki, G., 2025. Time, space, memory and brain-body rhythms. Nat. Rev. Neurosci. 10.1038/s41583-025-00987-2

Churchill, N.W., Spring, R., Grady, C., Cimprich, B., Askren, M.K., Reuter-Lorenz, P.A., Jung, M.S., Peltier, S., Strother, S.C., Berman, M.G., 2016. The suppression of scale-free fMRI brain dynamics across three different sources of effort: aging, task novelty and task difficulty. Sci. Rep. 6, 30895. 10.1038/srep30895

Coronel-Oliveros, C., Cofré, R., Orio, P., 2021. Cholinergic neuromodulation of inhibitory interneurons facilitates functional integration in whole-brain models. PLOS Comput. Biol. 17, e1008737. 10.1371/journal.pcbi.1008737

Coronel-Oliveros, C., Gießing, C., Medel, V., Cofré, R., Orio, P., 2023. Whole-brain modeling explains the context-dependent effects of cholinergic neuromodulation. NeuroImage 265, 119782. 10.1016/j.neuroimage.2022.119782

Cox, R., Rüber, T., Staresina, B.P., Fell, J., 2020. Phase-based coordination of hippocampal and neocortical oscillations during human sleep. Commun. Biol. 3, 176. 10.1038/s42003-020-0913-5

Das, A., Murphy, K., Drew, P.J., 2021. Rude mechanicals in brain haemodynamics: non-neural actors that influence blood flow. Philos. Trans. R. Soc. B Biol. Sci. 376, 20190635. 10.1098/rstb.2019.0635

David, O., Friston, K.J., 2003. A neural mass model for MEG/EEG: NeuroImage 20, 1743–1755. 10.1016/j.neuroimage.2003.07.015

Duff, E.P., Johnston, L.A., Xiong, J., Fox, P.T., Mareels, I., Egan, G.F., 2008. The power of spectral density analysis for mapping endogenous BOLD signal fluctuations. Hum. Brain Mapp. 29, 778–790. 10.1002/hbm.20601

Esteban, O., Markiewicz, C.J., Blair, R.W., Moodie, C.A., Isik, A.I., Erramuzpe, A., Kent, J.D., Goncalves, M., DuPre, E., Snyder, M., 2019. fMRIPrep: a robust preprocessing pipeline for functional MRI. Nat. Methods 16, 111–116.

Finn, E.S., Shen, X., Scheinost, D., Rosenberg, M.D., Huang, J., Chun, M.M., Papademetris, X., Constable, R.T., 2015. Functional connectome fingerprinting: identifying individuals using patterns of brain connectivity. Nat. Neurosci. 18, 1664–1671. 10.1038/nn.4135

Fox, M.D., Snyder, A.Z., Vincent, J.L., Corbetta, M., Van Essen, D.C., Raichle, M.E., 2005. The human brain is intrinsically organized into dynamic, anticorrelated functional networks. Proc. Natl. Acad. Sci. 102, 9673–9678. 10.1073/pnas.0504136102

Fox, M.D., Snyder, A.Z., Zacks, J.M., Raichle, M.E., 2006. Coherent spontaneous activity accounts for trial-to-trial variability in human evoked brain responses. Nat. Neurosci. 9, 23–25. 10.1038/nn1616

Friston, K.J., Holmes, A.P., Worsley, K.J., Poline, J. -P., Frith, C.D., Frackowiak, R.S.J., 1994. Statistical parametric maps in functional imaging: A general linear approach. Hum. Brain Mapp. 2, 189–210. 10.1002/hbm.460020402

Glasser, M.F., Coalson, T.S., Robinson, E.C., Hacker, C.D., Harwell, J., Yacoub, E., Ugurbil, K., Andersson, J., Beckmann, C.F., Jenkinson, M., Smith, S.M., Van Essen, D.C., 2016. A multi-modal parcellation of human cerebral cortex. Nature 536, 171–178. 10.1038/nature18933

Goldman, R.I., Stern, J.M., Engel, J., Cohen, M.S., 2002. Simultaneous EEG and fMRI of the alpha rhythm: NeuroReport 13, 2487–2492. 10.1097/00001756-200212200-00022

Golesorkhi, M., Gomez-Pilar, J., Tumati, S., Fraser, M., Northoff, G., 2021a. Temporal hierarchy of intrinsic neural timescales converges with spatial core-periphery organization. Commun. Biol. 4, 277. 10.1038/s42003-021-01785-z

Golesorkhi, M., Gomez-Pilar, J., Zilio, F., Berberian, N., Wolff, A., Yagoub, M.C.E., Northoff, G., 2021b. The brain and its time: intrinsic neural timescales are key for input processing. Commun. Biol. 4, 970. 10.1038/s42003-021-02483-6

Gonçalves, S.I., De Munck, J.C., Pouwels, P.J., Schoonhoven, R., Kuijer, J.P., Maurits, N.M., Hoogduin, J.M., Van Someren, E.J., Heethaar, R.M., Da Silva, F.L., 2006. Correlating the alpha rhythm to BOLD using simultaneous EEG/fMRI: inter-subject variability. Neuroimage 30, 203–213.

Gong, Z.-Q., Zuo, X.-N., 2025. Dark Brain Energy: Toward an Integrative Model of Spontaneous Slow Oscillations. Phys. Life Rev.

Grosenick, L., Klingenberg, B., Katovich, K., Knutson, B., Taylor, J.E., 2013. Interpretable whole-brain prediction analysis with GraphNet. NeuroImage 72, 304–321. 10.1016/j.neuroimage.2012.12.062

Gutierrez-Barragan, D., Basson, M.A., Panzeri, S., Gozzi, A., 2019. Infraslow State Fluctuations Govern Spontaneous fMRI Network Dynamics. Curr. Biol. 29, 2295–2306.e5. 10.1016/j.cub.2019.06.017

Harris, K.D., Thiele, A., 2011. Cortical state and attention. Nat. Rev. Neurosci. 12, 509–523. 10.1038/nrn3084

He, B.J., 2013. Spontaneous and Task-Evoked Brain Activity Negatively Interact. J. Neurosci. 33, 4672–4682. 10.1523/JNEUROSCI.2922-12.2013

He, B.J., 2011. Scale-Free Properties of the Functional Magnetic Resonance Imaging Signal during Rest and Task. J. Neurosci. 31, 13786–13795. 10.1523/JNEUROSCI.2111-11.2011

Huang, Z., Dai, R., Wu, Xuehai, Yang, Z., Liu, D., Hu, J., Gao, L., Tang, W., Mao, Y., Jin, Y., Wu, Xing, Liu, B., Zhang, Y., Lu, L., Laureys, S., Weng, X., Northoff, G., 2014. The self and its resting state in consciousness: An investigation of the vegetative state. Hum. Brain Mapp. 35, 1997–2008. 10.1002/hbm.22308

Huang, Z., Zhang, J., Longtin, A., Dumont, G., Duncan, N.W., Pokorny, J., Qin, P., Dai, R., Ferri, F., Weng, X., Northoff, G., 2017. Is There a Nonadditive Interaction Between Spontaneous and Evoked Activity? Phase-Dependence and Its Relation to the Temporal Structure of Scale-Free Brain Activity. Cereb. Cortex bhv288. 10.1093/cercor/bhv288

Jansen, B.H., Rit, V.G., 1995. Electroencephalogram and visual evoked potential generation in a mathematical model of coupled cortical columns. Biol. Cybern. 73, 357–366. 10.1007/BF00199471

Ji, J.L., Spronk, M., Kulkarni, K., Repovš, G., Anticevic, A., Cole, M.W., 2019. Mapping the human brain’s cortical-subcortical functional network organization. NeuroImage 185, 35–57. 10.1016/j.neuroimage.2018.10.006

Kasagi, M., Huang, Z., Narita, K., Shitara, H., Motegi, T., Suzuki, Y., Fujihara, K., Tanabe, S., Kosaka, H., Ujita, K., 2017. Association between scale-free brain dynamics and behavioral performance: Functional MRI study in resting state and face processing task. Behav. Neurol. 2017, 2824615.

Klar, P., Çatal, Y., Langner, R., Huang, Z., Northoff, G., 2023a. Scale-free dynamics in the core-periphery topography and task alignment decline from conscious to unconscious states. Commun. Biol. 6, 499. 10.1038/s42003-023-04879-y

Klar, P., Çatal, Y., Langner, R., Huang, Z., Northoff, G., 2023b. Scale-free dynamics of core-periphery topography. Hum. Brain Mapp. 44, 1997–2017. 10.1002/hbm.26187

Lakatos, P., Gross, J., Thut, G., 2019. A new unifying account of the roles of neuronal entrainment. Curr. Biol. 29, R890–R905.

Lakatos, P., Karmos, G., Mehta, A.D., Ulbert, I., Schroeder, C.E., 2008. Entrainment of Neuronal Oscillations as a Mechanism of Attentional Selection. Science 320, 110–113. 10.1126/science.1154735

Li, M., Han, Y., Aburn, M.J., Breakspear, M., Poldrack, R.A., Shine, J.M., Lizier, J.T., 2019. Transitions in information processing dynamics at the whole-brain network level are driven by alterations in neural gain. PLOS Comput. Biol. 15, e1006957. 10.1371/journal.pcbi.1006957

Liu, D.-Y., Li, M., Yu, J., Gao, Y., Zhang, X., Hu, D., Northoff, G., Song, X.M., Zhu, J., 2024. Sex differences in the human brain related to visual motion perception. Biol. Sex Differ. 15, 92. 10.1186/s13293-024-00668-2

Lu, F.-M., Wang, Y.-F., Zhang, J., Chen, H.-F., Yuan, Z., 2017. Optical mapping of the dominant frequency of brain signal oscillations in motor systems. Sci. Rep. 7, 14703. 10.1038/s41598-017-15046-9

Maris, E., Oostenveld, R., 2007. Nonparametric statistical testing of EEG- and MEG-data. J. Neurosci. Methods 164, 177–190. 10.1016/j.jneumeth.2007.03.024

Moran, R.J., Kiebel, S.J., Stephan, K.E., Reilly, R.B., Daunizeau, J., Friston, K.J., 2007. A neural mass model of spectral responses in electrophysiology. NeuroImage 37, 706–720. 10.1016/j.neuroimage.2007.05.032

Murphy, K., Fox, M.D., 2017. Towards a consensus regarding global signal regression for resting state functional connectivity MRI. NeuroImage 154, 169–173. 10.1016/j.neuroimage.2016.11.052

Nau, M., Schmid, A.C., Kaplan, S.M., Baker, C.I., Kravitz, D.J., 2024. Centering cognitive neuroscience on task demands and generalization. Nat. Neurosci. 27, 1656–1667. 10.1038/s41593-024-01711-6

Nichols, T.E., Holmes, A.P., 2002. Nonparametric permutation tests for functional neuroimaging: A primer with examples. Hum. Brain Mapp. 15, 1–25. 10.1002/hbm.1058

Norcia, A.M., Appelbaum, L.G., Ales, J.M., Cottereau, B.R., Rossion, B., 2015. The steady-state visual evoked potential in vision research: A review. J. Vis. 15, 4. 10.1167/15.6.4

Northoff, G., Buccellato, A., Zilio, F., 2024. Connecting brain and mind through temporo-spatial dynamics: Towards a theory of common currency. Phys. Life Rev. 52, 29–43.

Palva, J.M., Palva, S., 2012. Infra-slow fluctuations in electrophysiological recordings, blood-oxygenation-level-dependent signals, and psychophysical time series. Neuroimage 62, 2201–2211.

Palva, S., Palva, J.M., 2018. Roles of Brain Criticality and Multiscale Oscillations in Temporal Predictions for Sensorimotor Processing. Trends Neurosci. 41, 729–743. 10.1016/j.tins.2018.08.008

Raichle, M.E., Mintun, M.A., 2006. BRAIN WORK AND BRAIN IMAGING. Annu. Rev. Neurosci. 29, 449–476. 10.1146/annurev.neuro.29.051605.112819

Reynolds, J.H., Heeger, D.J., 2009. The Normalization Model of Attention. Neuron 61, 168–185. 10.1016/j.neuron.2009.01.002

Rosen, B.Q., Halgren, E., 2021. A Whole-Cortex Probabilistic Diffusion Tractography Connectome. eneuro 8, ENEURO.0416-20.2020. 10.1523/ENEURO.0416-20.2020

Sasai, S., Koike, T., Sugawara, S.K., Hamano, Y.H., Sumiya, M., Okazaki, S., Takahashi, H.K., Taga, G., Sadato, N., 2021. Frequency-specific task modulation of human brain functional networks: A fast fMRI study. Neuroimage 224, 117375. 10.1016/j.neuroimage.2020.117375

Scheeringa, R., Fries, P., Petersson, K.-M., Oostenveld, R., Grothe, I., Norris, D.G., Hagoort, P., Bastiaansen, M.C.M., 2011. Neuronal Dynamics Underlying High- and Low-Frequency EEG Oscillations Contribute Independently to the Human BOLD Signal. Neuron 69, 572–583. 10.1016/j.neuron.2010.11.044

Schroeder, C.E., Lakatos, P., 2009. Low-frequency neuronal oscillations as instruments of sensory selection. Trends Neurosci. 32, 9–18. 10.1016/j.tins.2008.09.012

Servan-Schreiber, D., Printz, H., Cohen, J.D., 1990. A Network Model of Catecholamine Effects: Gain, Signal-to-Noise Ratio, and Behavior. Science 249, 892–895. 10.1126/science.2392679

Shine, J.M., Aburn, M.J., Breakspear, M., Poldrack, R.A., 2018. The modulation of neural gain facilitates a transition between functional segregation and integration in the brain. eLife 7, e31130. 10.7554/eLife.31130

Shulman, G.L., Fiez, J.A., Corbetta, M., Buckner, R.L., Miezin, F.M., Raichle, M.E., Petersen, S.E., 1997. Common Blood Flow Changes across Visual Tasks: II. Decreases in Cerebral Cortex. J. Cogn. Neurosci. 9, 648–663. 10.1162/jocn.1997.9.5.648

Thiele, A., Bellgrove, M.A., 2018. Neuromodulation of Attention. Neuron 97, 769–785. 10.1016/j.neuron.2018.01.008

Toffanin, P., De Jong, R., Johnson, A., Martens, S., 2009. Using frequency tagging to quantify attentional deployment in a visual divided attention task. Int. J. Psychophysiol. 72, 289–298. 10.1016/j.ijpsycho.2009.01.006

Tomasi, D., Wang, G.-J., Volkow, N.D., 2013. Energetic cost of brain functional connectivity. Proc. Natl. Acad. Sci. 110, 13642–13647. 10.1073/pnas.1303346110

Uddin, L.Q., 2017. Mixed Signals: On Separating Brain Signal from Noise. Trends Cogn. Sci. 21, 405–406. 10.1016/j.tics.2017.04.002

Van Atteveldt, N., Musacchia, G., Zion-Golumbic, E., Sehatpour, P., Javitt, D.C., Schroeder, C., 2015. Complementary fMRI and EEG evidence for more efficient neural processing of rhythmic vs. unpredictably timed sounds. Front. Psychol. 6, 1663.

Vialatte, F.-B., Maurice, M., Dauwels, J., Cichocki, A., 2010. Steady-state visually evoked potentials: Focus on essential paradigms and future perspectives. Prog. Neurobiol. 90, 418–438. 10.1016/j.pneurobio.2009.11.005

Wang, Y., Chen, W., Ye, L., Biswal, B.B., Yang, X., Zou, Q., Yang, P., Yang, Q., Wang, X., Cui, Q., Duan, X., Liao, W., Chen, H., 2018. Multiscale energy reallocation during low-frequency steady-state brain response. Hum. Brain Mapp. 39, 2121–2132. 10.1002/hbm.23992

Wang, Y., Long, Z., Cui, Q., Liu, F., Jing, X.-J., Chen, H., Guo, X.-N., Yan, J.H., Chen, H.-F., 2016. Low frequency steady-state brain responses modulate large scale functional networks in a frequency-specific means. Hum. Brain Mapp. 37, 381–394.

Wang, Y.-F., Dai, G.-S., Liu, F., Long, Z.-L., Yan, J.H., Chen, H.-F., 2015. Steady-state BOLD Response to Higher-order Cognition Modulates Low-Frequency Neural Oscillations. J. Cogn. Neurosci. 27, 2406–2415. 10.1162/jocn_a_00864

Wang, Y.-F., Liu, F., Long, Z.-L., Duan, X.-J., Cui, Q., Yan, J.H., Chen, H.-F., 2014. Steady-State BOLD Response Modulates Low Frequency Neural Oscillations. Sci. Rep. 4, 7376. 10.1038/srep07376

Waschke, L., Kloosterman, N.A., Obleser, J., Garrett, D.D., 2021. Behavior needs neural variability. Neuron 109, 751–766. 10.1016/j.neuron.2021.01.023

Weissman, D.H., Roberts, K.C., Visscher, K.M., Woldorff, M.G., 2006. The neural bases of momentary lapses in attention. Nat. Neurosci. 9, 971–978. 10.1038/nn1727

Wieser, M.J., Miskovic, V., Keil, A., 2016. Steady-state visual evoked potentials as a research tool in social affective neuroscience. Psychophysiology 53, 1763–1775. 10.1111/psyp.12768

Wolff, A., Berberian, N., Golesorkhi, M., Gomez-Pilar, J., Zilio, F., Northoff, G., 2022. Intrinsic neural timescales: temporal integration and segregation. Trends Cogn. Sci. 26, 159–173. 10.1016/j.tics.2021.11.007

Wolman, A., Çatal, Y., Klar, P., Steffener, J., Northoff, G., 2024. Repertoire of timescales in uni - and transmodal regions mediate working memory capacity. NeuroImage 291, 120602. 10.1016/j.neuroimage.2024.120602

Yan, C.G., Wang, X.D., Zuo, X.N., Zang, Y.F., 2016. DPABI: Data Processing & Analysis for (Resting-State) Brain Imaging. Neuroinformatics 14, 339–51. 10.1007/s12021-016-9299-4

Zhang, C., Wang, Y., Jing, X., Yan, J.H., 2023. Brain mechanisms of mental processing: from evoked and spontaneous brain activities to enactive brain activity. Psychoradiology 3, kkad010. 10.1093/psyrad/kkad010

Zhang, S., Li, C.-S.R., 2012. Task-Related, Low-Frequency Task-Residual, and Resting State Activity in the Default Mode Network Brain Regions. Front. Psychol. 3. 10.3389/fpsyg.2012.00172

Zou, H., Hastie, T., 2005. Addendum: Regularization and Variable Selection Via the Elastic Net. J. R. Stat. Soc. Ser. B Stat. Methodol. 67, 768–768. 10.1111/j.1467-9868.2005.00527.x

