## Supplementary materials file for "Temporal structure of task engagement reorganizes infra-slow BOLD dynamics"

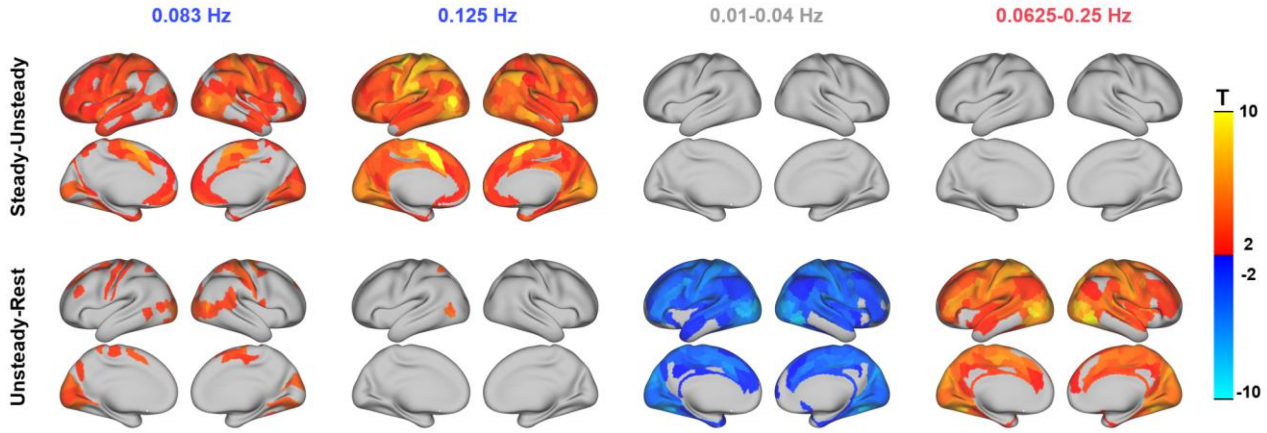


**Supplementary Figure 1. Cortical maps of task-related spectral power differences across frequency ranges of interest.** Region-wise paired-sample t-statistics comparing fractional BOLD PSD between conditions in the UESTC dataset. Rows indicate condition contrasts: steady-unsteady (top) and unsteady-rest (bottom). Columns indicate frequency ranges: the stimulus-locked frequencies of 0.083 Hz and 0.125 Hz, the spontaneous low infra-slow range of 0.01-0.04 Hz, and the broader task-relevant range of 0.0625-0.25 Hz.

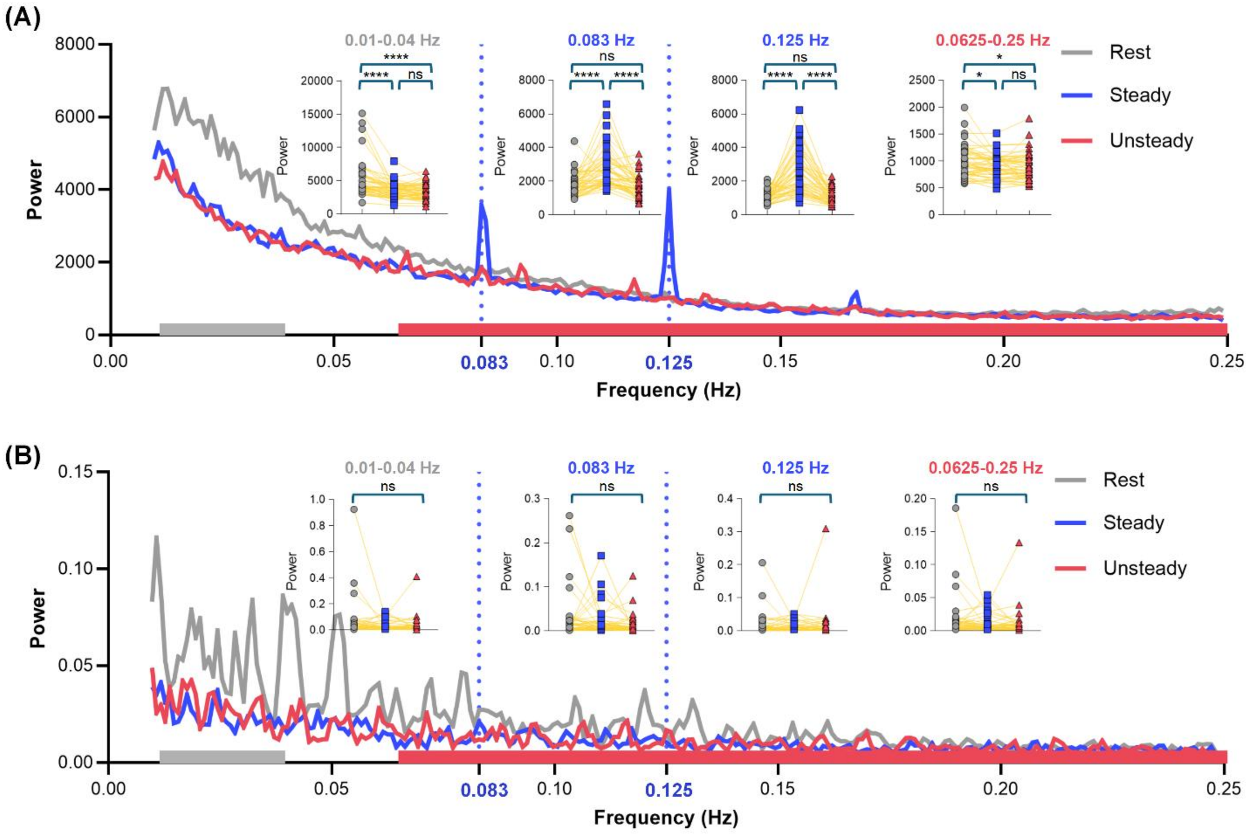


**Supplementary Figure 2. Raw (non-normalized) power spectral density and head motion spectral analysis. (A)** Whole-brain averaged raw PSD (without normalization) of BOLD signals for resting (gray), steady-state (blue), and unsteady-state (red) conditions in the UESTC dataset. The stimulus-locked spectral peaks at 0.083 Hz and 0.125 Hz remain clearly visible in the steady-state condition, confirming that the power redistribution effect is not an artifact of the normalization procedure. Insets show condition comparisons at four frequency bands. **(B)** PSD of mean framewise displacement (FD), a summary measure of head motion, across the three conditions. No significant differences are observed between conditions at any frequency band (all ns), indicating that the spectral peaks in the BOLD signal are not driven by condition-related differences in head motion.

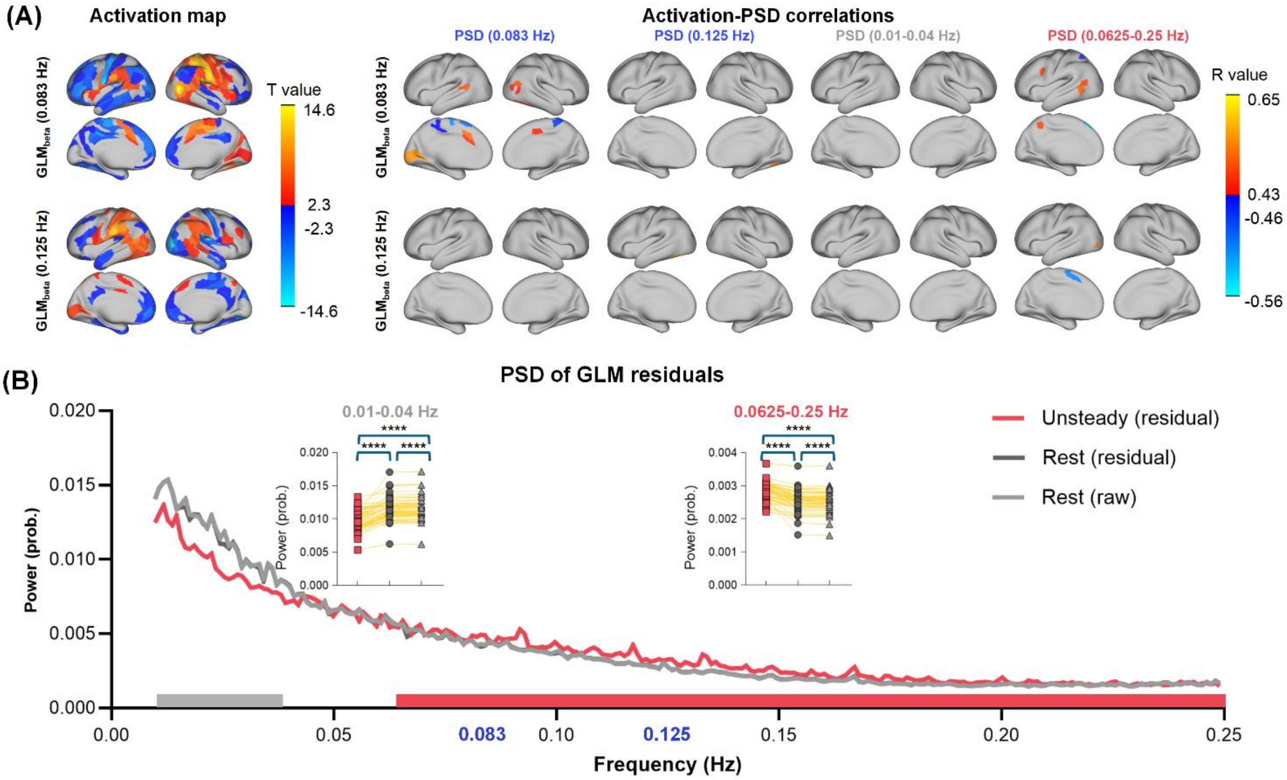


**Supplementary Figure 3. GLM activation and spectral analysis for the unsteady-state condition.** (A) Left: Cortical t-statistic maps of GLM-derived activation in the unsteady-state condition for the two stimulus-locked frequencies, 0.083 Hz and 0.125 Hz. Right: For each cortical parcel, Pearson correlations were computed across subjects between GLM beta values and PSD power, and the resulting correlation coefficients were displayed on the cortical surface. Rows indicate GLM beta maps for the 0.083-Hz and 0.125-Hz regressors; columns indicate PSD power at 0.083 Hz, 0.125 Hz, 0.01-0.04 Hz, and 0.0625-0.25 Hz. (B) Whole-brain averaged PSD of steady-state GLM residuals, compared with raw resting-state PSD and procedure-matched resting-state GLM residuals at 0.01-0.04 Hz and 0.0625-0.25 Hz.

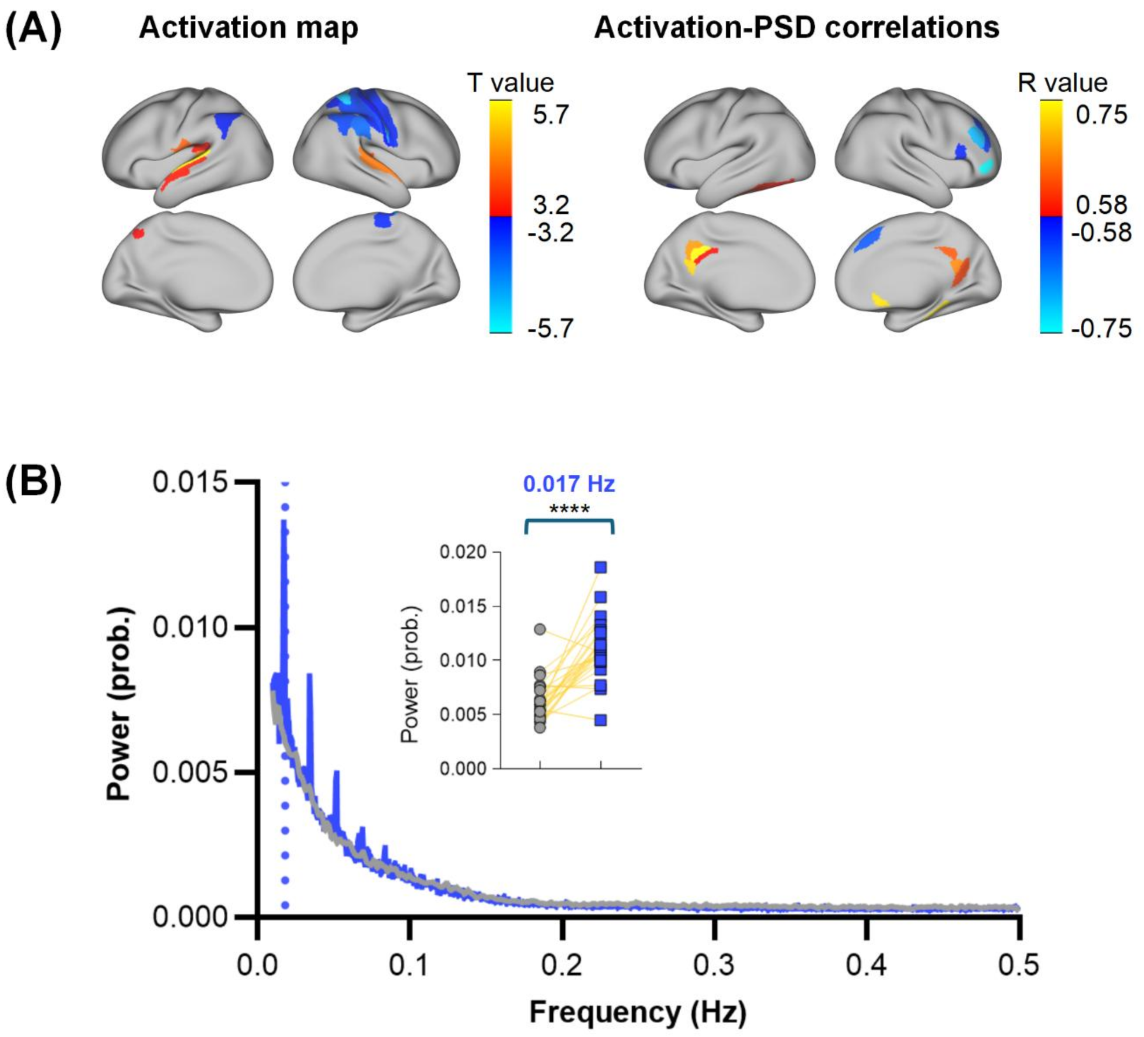


**Supplementary Figure 4. GLM activation and spectral analysis in HZNU dataset.** (A) Left: Cortical t-statistic maps of GLM-derived activation in task state. Right: Pearson correlations were computed across subjects between GLM beta values and PSD power, and the resulting correlation coefficients were displayed on the cortical surface. (B) Whole-brain averaged PSD of steady-state GLM residuals, compared with raw resting-state PSD and procedure-matched resting-state GLM residuals at 0.017 Hz.

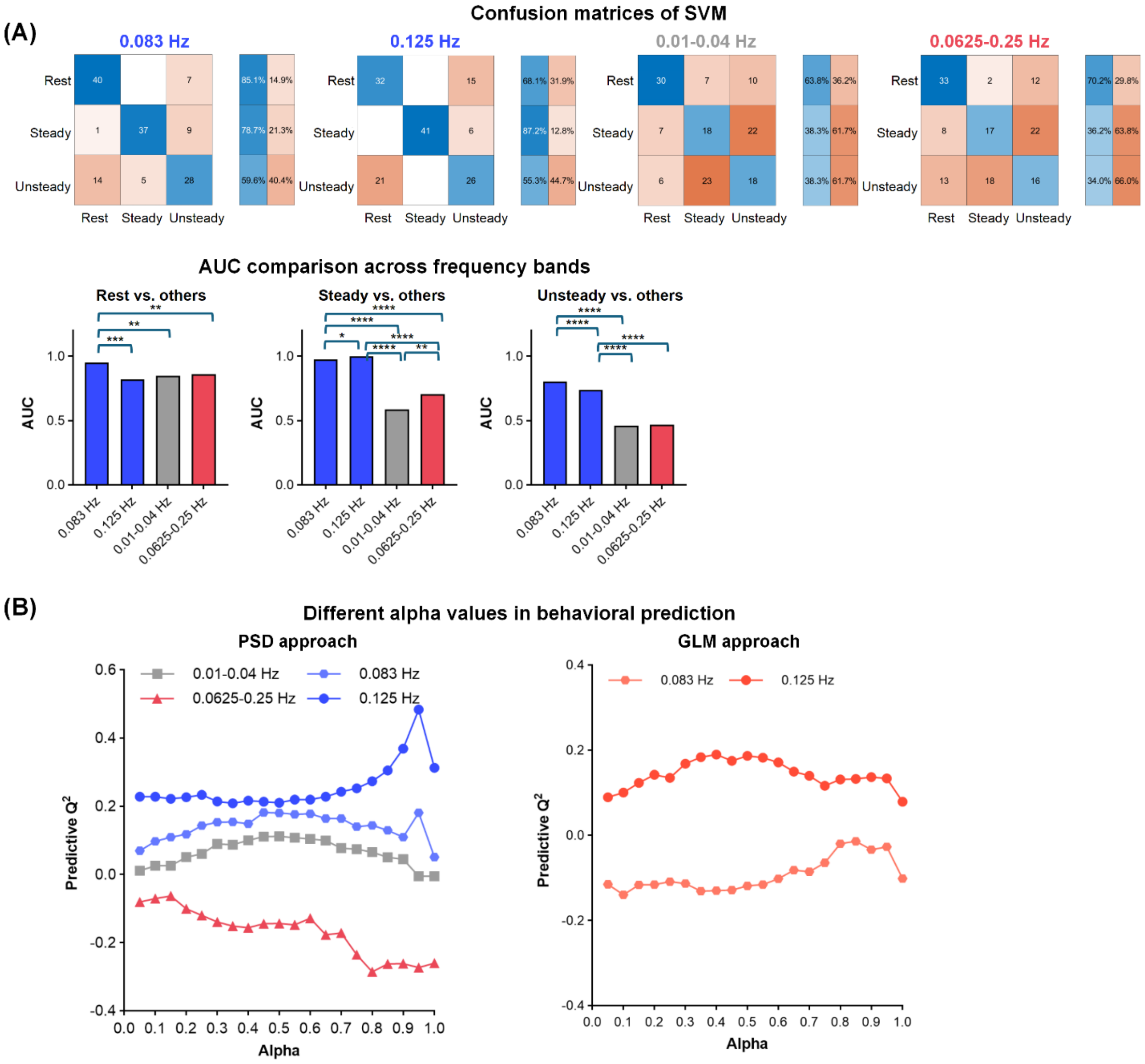


**Supplementary Figure 5. Classification and behavioral prediction based on PSD features in the UESTC dataset. (A)** Top: Multiclass classification of Rest, Steady, and Unsteady states using PSD features at the stimulation frequencies, 0.083 Hz and 0.125 Hz, and two broader frequency ranges, 0.01-0.04 Hz and 0.0625-0.25 Hz. Rows indicate true labels and columns indicate predicted labels; adjacent bars show condition-wise accuracy and error rate. Bottom: One-versus-rest AUC for each frequency band and condition, with pairwise DeLong tests comparing classification performance across bands. **(B)** Predictive $Q^2$ of elastic net models as a function of the L1/L2 mixing parameter for PSD features and GLM-derived activation features. The 0.125-Hz PSD model showed the strongest and most stable behavioral prediction across the tested mixing-parameter range. Results were based on leave-one-out cross-validation.

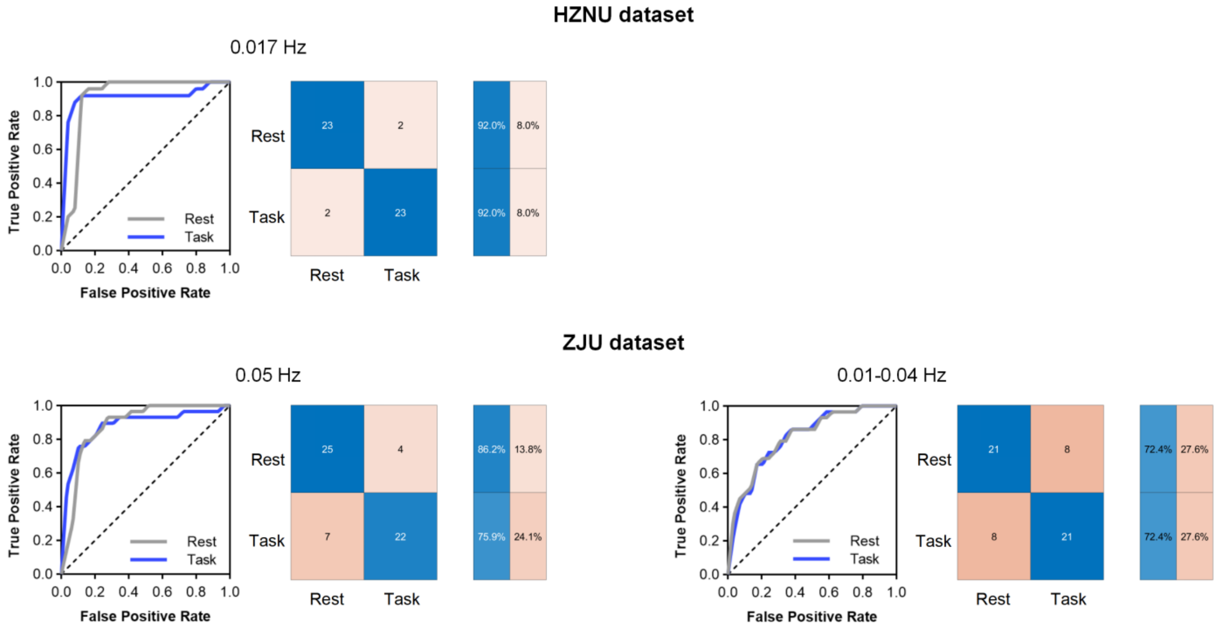


**Supplementary Figure 6. Replication of PSD-based rest-task classification in independent datasets.** Rest-task classification based on PSD features was replicated in the HZNU and ZJU datasets using dataset-specific stimulation frequencies. In the HZNU dataset, classification based on 0.017 Hz PSD achieved high discrimination for both Rest and Task conditions, with AUCs of AUC_Rest = 0.93 and AUC_Task = 0.93, respectively. In the ZJU dataset, classification based on 0.05-Hz PSD also showed robust performance, with AUCs of AUC_Rest = 0.89 and AUC_Task = 0.90. The low-frequency comparison band in the ZJU dataset, 0.01-0.04 Hz, yielded AUCs of AUC_Rest = 0.83 and AUC_Task = 0.833. ROC curves, confusion matrices, and condition-wise accuracy/error bars show consistent rest-task discrimination across independent datasets.

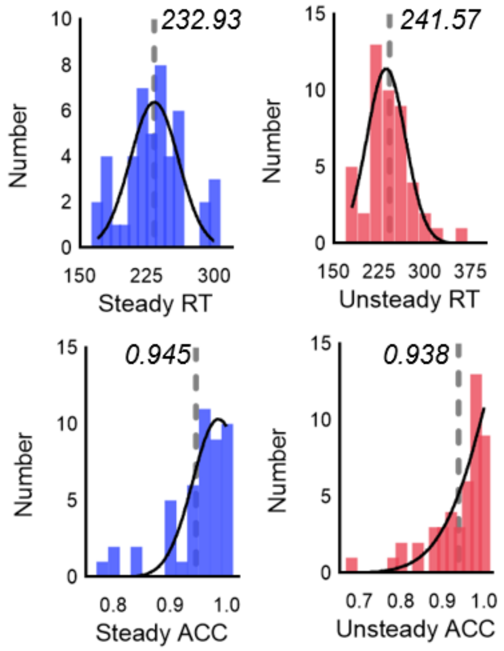

**Supplementary Figure 7. Distribution of behavioral performance measures.** Histograms of mean reaction time (RT; top) and accuracy (ACC; bottom) for steady-state (blue, left) and unsteady-state (red, right) conditions. Black curves indicate fitted distributions. Dashed lines and values denote group means (steady-state: mean RT = 232.93 ms, mean ACC = 0.945; unsteady-state: mean RT = 241.57 ms, mean ACC = 0.938).

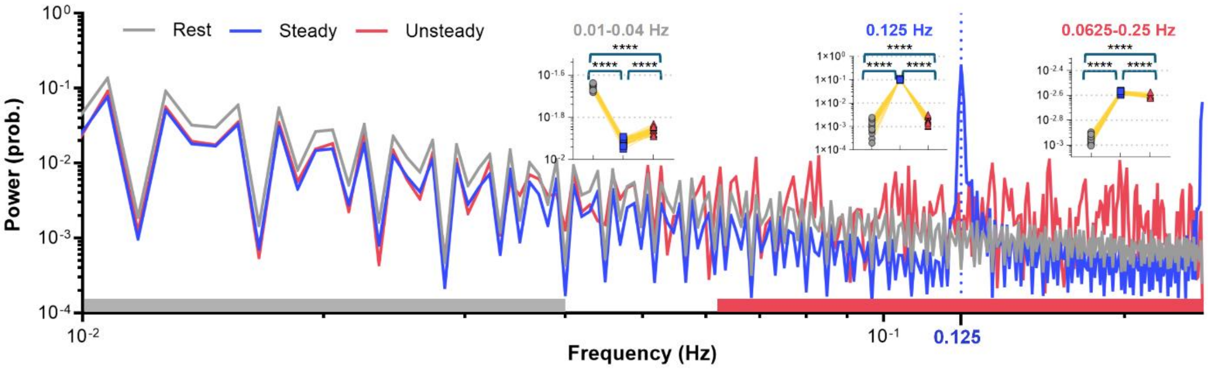


**Supplementary Figure 8.** Power spectral density of simulated BOLD signals generated by the Jansen-Rit neural mass model. Single-node BOLD power spectral density (PSD) is shown for rest, steady-state, and unsteady-state conditions. Colored bars indicate the frequency ranges of interest: 0.01-0.04 Hz, the stimulus-locked frequencies of 0.083 Hz and 0.125 Hz, and the broader task-relevant range of 0.0625-0.25 Hz. Insets show paired subject-level comparisons.
